# Cell-specific alterations in *Pitx1* regulatory landscape activation caused by the loss of a single enhancer

**DOI:** 10.1101/2021.03.10.434611

**Authors:** Raquel Rouco, Olimpia Bompadre, Antonella Rauseo, Olivier Fazio, Fabrizio Thorel, Rodrigue Peraldi, Guillaume Andrey

## Abstract

Most developmental genes rely on multiple transcriptional enhancers for their accurate expression during embryogenesis. Because enhancers may have partially redundant activities, the loss of one of them often leads to a partial loss of gene expression and concurrent moderate phenotypic outcome, if any. While such a phenomenon has been observed in many instances, the nature of the underlying mechanisms remains elusive. We used the *Pitx1* testbed locus to characterize in detail the regulatory and cellular identity alterations following the deletion *in vivo* of one of its enhancers (*Pen*), which normally accounts for 30 percent of *Pitx1* expression in hindlimb buds. By combining single cell transcriptomics and a novel *in embryo* cell tracing approach, we observed that this global decrease in *Pitx1* expression results from both an increase in the number of non- or low-expressing cells, and a decrease in the number of high-expressing cells. We found that the over-representation of *Pitx1* non/low-expressing cells originates from a failure of the *Pitx1* locus to coordinate enhancer activities and 3D chromatin changes. The resulting increase in *Pitx1* non/low-expressing cells eventually affects the proximal limb more severely than the distal limb, leading to a clubfoot phenotype likely produced through a localized heterochrony and concurrent loss of irregular connective tissue. This data suggests that, in some cases, redundant enhancers may be used to locally enforce a robust activation of their host regulatory landscapes.

## Introduction

Alteration in the enhancer composition of regulatory landscapes at developmental genes can lead to pathologies by modifying the dosage and/or distribution of genes transcription (Kvon et al. 2021). Indeed, over the past years, loss of single regulatory units within complex and partially redundant regulatory landscapes were shown to have clear phenotypical outcomes despite inducing only partial decreases in average transcription (Montavon et al. 2011; Will et al. 2017; Osterwalder et al. 2018). As the alterations in the regulatory mechanisms following enhancer deletion have mostly been characterized using bulk tissue analysis, it has been difficult to determine the cell-specific variability behind the loss of expression that accounts for phenotypes. In order to understand the precise molecular origin of these phenotypes, it is therefore essential to characterize how a single enhancer contributes to the activation of entire regulatory landscapes in specific cell populations. An effective model system to address these unsolved questions is the limb bud, where organogenesis requires a tight control of gene transcription to achieve correct patterning (Petit et al. 2017). Critical to this is *Pitx1*, a transcription factor coding gene that is normally expressed in developing hindlimb buds, but not in forelimbs, which channels the limb development program to differentiate into a leg (Lanctot et al. 1997; Infante et al. 2013; Nemec et al. 2017). Consequently, forelimb *Pitx1* gain-of-function can induce an arm-to-leg transformation, featured by the appearance of an ectopic patella as well as complex changes in the muscular and tendon wiring (DeLaurier et al. 2006; Kragesteen et al. 2018). In contrast, *Pitx1* knock out has been shown to induce partial leg-to-arm transformations with the disappearance of the patella as well as long bone dysplasia and polydactyly (Lanctot et al. 1999b; DeLaurier et al. 2006). Unexpectedly, bulk transcriptomics strategies have only revealed marginal downstream gene expression changes upon *Pitx1* loss, suggesting that an interplay between these changes and the growth rate of limb cell subpopulations collectively result in the various phenotypes (Lanctot et al. 1999b; DeLaurier et al. 2006; Alvarado et al. 2011; Nemec et al. 2017; Kragesteen et al. 2018).

As for many developmental genes, several enhancers coordinate *Pitx1* expression in hindlimbs, among which is *Pen*, previously shown to account for 35-50% of *Pitx1* hindlimb expression (Kragesteen et al. 2018; Thompson et al. 2018). The deletion of *Pen* has no impact on bone length or digit numbers but induces a partially penetrant clubfoot phenotype, similar to the one observed in mice and humans upon *Pitx1* haploinsufficiency (Alvarado et al. 2011; Kragesteen et al. 2018). One particularity to the *Pitx1* locus is that it establishes fundamentally different 3D chromatin conformation in transcriptionally active hindlimbs and inactive forelimbs. In active hindlimbs, *Pitx1* forms chromatin interactions with cognate *cis*-regulatory regions spread over 400 kbs, including *Pen* as well as *Pit, RA4* and *PelB*. In contrast, in inactive forelimbs these interactions are absent and the *Pitx1* gene forms a contact with the polycomb-repressed gene *Neurog1* (Kragesteen et al. 2018). In this work, we use a combination of single cell transcriptomics (scRNA-seq), a fluorescent cell-tracing approach and genomic technologies to define the contribution of a single enhancer (*Pen*) in establishing the epigenetically- and structurally-active *Pitx1* regulatory landscape. Moreover, we investigate whether changes in enhancer activities or 3D structure fundamentally associate with transcription or if those can be functionally disconnected of the transcriptional process. Finally, we assessed if *Pitx1* expression is homogenous across limb cell population and if distinct expression levels rely on different enhancer repertoires or, alternatively, in progressive changes in cis-regulatory landscape activities.

## Results

### Two approaches to track *Pitx1* activities suggest a bimodal *cis*-regulatory behavior

In order to characterize transcriptional, chromatin and structural changes following the *Pen* enhancer deletion, we combined genetic manipulation of the *Pitx1* locus with scRNA-seq and chromatin analysis of sorted limb cell populations. Both approaches enable characterization of complementary features of gene transcriptional regulation following alterations of the cis-regulatory landscape.

First, to define the hindlimb cell types that are expressing *Pitx1* and to assess how the *Pen* enhancer regulates its expression in these cells, we generated single-cell preparations from wildtype fore- and hindlimb buds as well as *Pen* enhancer deleted (*Pitx1*^*Pen-/Pen-*^) or *Pitx1* knocked-out (*Pitx1*^*-/-*^) hindlimbs (**Fig. 1A**). We performed 10x genomics in duplicates from E12.5 limb buds as these correspond to a transition stage between patterning and cell-differentiation phases. By performing unsupervised clustering of all the wildtype and mutant single cell transcriptomic datasets, we identified five clusters (**Fig 1B**) corresponding to the main populations of the limb: one mesenchymal cluster (*Prrx1+*; 90% of the cells) and four satellite clusters including muscle (*Myod1+, Ttn+*; 4% of the cells), epithelium (*Wnt6+, Krt17+*; 5% of the cells), endothelium (*Cdh5+, Cldn5+*; 1.5% of the cells) and one immune cell cluster (*C1qa, Ccr1+*; 0.05% of the cells) (**Supplementary Table S1**). Yet, as *Pitx1* is expressed only in the hindlimb mesenchymal cluster (**Fig 1C**), further analyses will be performed only in these cells.

**Figure 1.**
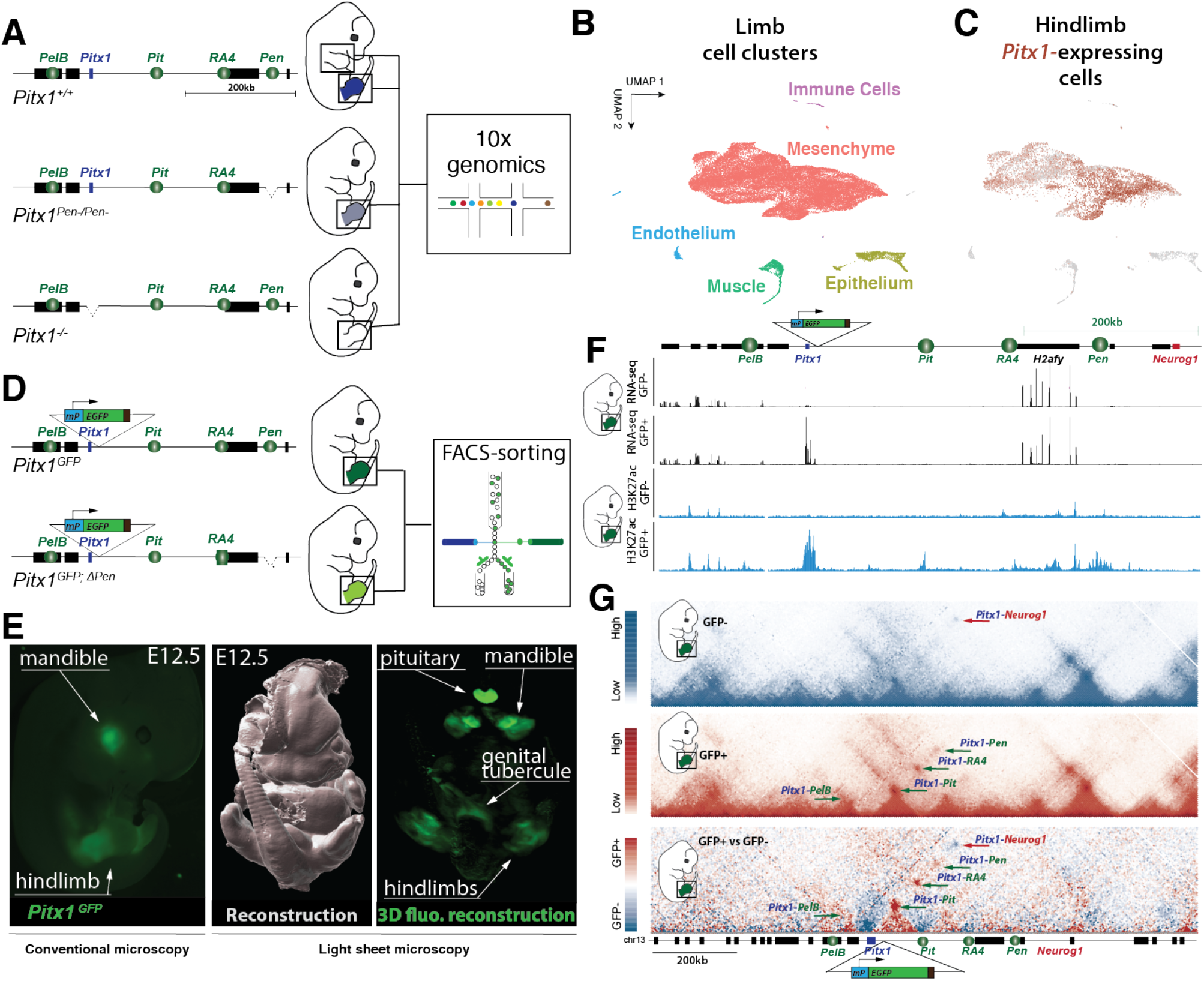
Experimental setup, single cell clustering and regulatory sensor. **A**. Wildtype, *Pitx1*^*Pen-/Pen*-^ and *Pitx1*^*-/-*^transgenic E12.5 embryos were obtained by tetraploid complementation and single cell transcriptomic analyses were produced from fore- and hindlimbs. **B**. UMAP clustering of wildtype and mutant fore- and hindlimbs shows one mesenchymal as well as four satellite clusters. **C**. UMAP colored according to *Pitx1* expression shows expression only in the mesenchyme cluster. **D**. A cassette containing a minimal *β-globin* promoter (*mP*) and an EGFP reporter gene is integrated upstream of *Pitx1*. A secondary CRISPR/Cas9 targeting is then used to delete the *Pen* enhancer. **E**. Conventional and light sheet microscopy reveal that *Pitx1*^*GFP*^ embryos display EGFP expression domains corresponding to the one of *Pitx1*. **F**. RNA-seq and H3K27ac of sorted hindlimb cells show that the sensor approach can separate *Pitx1* active (GFP+) and inactive (GFP-) regulatory landscapes. **G**. The 3D structure of active and inactive landscapes in the hindlimbs is fundamentally different. GFP+ cells bear chromatin interactions between *Pitx1* and its associated enhancers. GFP-cells do not display these interactions but a strong contact between *Pitx1* and *Neurog1*.

In parallel, we devised a fluorescent reporter system to track the regulatory activities of the *Pitx1* locus in hindlimbs (**Fig 1D**). Specifically, we first established a reporter line (*Pitx1*^*GFP*^) by homozygously integrating a regulatory sensor cassette, constituted of a minimal *β-globin* promoter and an EGFP reporter gene, 2kb upstream of the *Pitx1* promoter in mouse embryonic stem cells (mESCs). These cells were then re-targeted to obtain a homozygous deletion of the *Pen* enhancer (*Pitx1*^*GFP;ΔPen*^). Embryos were then derived from the mESCs *via* tetraploid complementation (Artus and Hadjantonakis 2011). Conventional and light sheet imaging of *Pitx1*^*GFP*^embryos showed that the reporter was expressed in all *Pitx1* expression domains including the pituitary gland, the mandible, the genital tubercle and the hindlimbs (**Fig. 1D, S1A, Supplementary Video S1)** (Lanctot et al. 1999a; Lanctot et al. 1999b; Chiu et al. 2010).

We then FACS sorted GFP+ and GFP-cells from E12.5 hindlimbs and processed cells for RNA-seq, ChIP-seq and Capture-HiC (C-HiC) (**Fig. 1F-G, S1B-C**). We found that 8% of the cells in wildtype hindlimbs displayed no EGFP signal, thereby suggesting that a majority of hindlimb cells possess an active *Pitx1* regulatory landscape. Following cell sorting, we next compared the transcriptome of GFP+ and GFP-cells and observed a 35-fold enrichment for *Pitx1* expression in GFP+ cells, validating the *Pitx1*^*GFP*^ allele to track the *Pitx1* regulatory landscape activities (**Fig. 1F, S2A, Supplementary Table S2**). As expected from our scRNA-seq analyses, we found that GFP+/*Pitx1*+ cells were enriched for limb mesenchymal derivatives markers (*Prrx1, Sox9, Col2a1, Col3a1, Lum*) and that GFP-/*Pitx1*-were enriched for satellite cluster markers including muscle (*Myod1, Ttn*), epithelium (*Wnt6, Krt17*), endothelium (*Cdh5, Cldn5*) and immune cells (*C1qa, Ccr1*) (**Fig. S2B, Supplementary Table S2**). Yet, the enrichment of these cell types does not preclude a fraction of GFP-/*Pitx1*-to be of mesenchymal origin as we found a weak but clear expression of some mesenchymal markers such as *Prrx1* or *Twist1* in this population (**Fig. S2C)**

We then assayed the *cis*-regulatory activities in GFP-/Pitx1- and GFP+/Pitx1+ hindlimb cells using the H3K27ac chromatin mark as a proxy for enhancer activities and C-HiC to determine the locus chromatin architecture (Rada-Iglesias et al. 2011). In GFP-/Pitx1-cells, neither *Pitx1* promoter nor its various enhancers, including *Pen*, were found enriched with H3K27ac (**Fig. 1E, 1F**). Moreover, the locus 3D structure is in a repressed state where *Pitx1* displays a strong interaction with the repressed *Neurog1* gene and no interaction with its cognate enhancers. This data show that GFP-/Pitx1-hindlimb cells display a complete absence of active regulatory landscape features. In contrast, in GFP+/Pitx1+ cells all known *Pitx1* enhancers as well as its promoter are strongly enriched in H3K27ac chromatin marks. Furthermore, in these cells *Pitx1* establishes strong contact with its cognate enhancers *PelB, Pit, RA4* and *Pen* (**Fig. 1E, 1F**).

In summary, this data shows that within the hindlimb, classically considered as a *Pitx1* active tissue, 8% of cells, from mesenchymal, immune, endothelium, muscle and epithelium origin, display an inactive *Pitx1 cis*-regulatory landscape and 3D architecture. Moreover, it suggests a *bimodal* regulatory behavior, where the *Pitx1* promoter, its associated enhancers and the locus 3D structure are all displaying an *active* mode or none of them are. We then further characterize *Pitx1* expression specificities within the hindlimb mesenchyme.

### Hindlimb proximal cell clusters express *Pitx1* at higher level

To characterize *Pitx1* transcription within mesenchymal subpopulations, we first re-clustered mesenchymal cells from all datasets. From this analysis, we could define nine clusters (**Fig. 2A**). We first observed that their distribution in the UMAP space is strongly influenced by the limb proximo-distal axis, as illustrated by *Shox2* (proximal marker) and *Hoxd13* (distal marker) transcript distribution (**Fig. 2B**). We further annotated the clusters according to the expression of known marker genes. In the proximal limb section, we identified four clusters. First, we found an undifferentiated **P**roximal **P**roliferative **P**rogenitors cluster which is characterized by high expression of proliferative marker genes (**PPP**: *Shox2+, Hist1h1d*+, *Top2a+*). We then identified a **T**endon **P**rogenitor cluster (**TP**: *Shox2+*; *Osr1+; Scx+*) and an **I**rregular **C**onnective **T**issue cluster which includes muscle connective tissue and ultimately patterns tendons and muscles (**ICT**: *Shox2*+; *Osr1+, Dcn+, Lum+, Kera+, Col3a1+*)(Besse et al. 2020). Finally, in the proximal limb we observed a single cluster of **P**roximal **C**ondensations that will give rise to proximal limb bones (**PC**: *Tbx15+; Sox9+*; *Col9a3+*). In the distal limb, we observed the presence of two undifferentiated distal mesenchyme (*Msx1*+) clusters: one that we classified as **D**istal **P**roliferative **P**rogenitors (**DPP**: *Hoxd13+*; *Msx1+*; *Hist1h1d+*) as it displays a strong expression of proliferation markers, while the other is defined as **D**istal **P**rogenitors (**DP**: *Hoxd13*+; *Msx1*+). Also, in the distal limb, we identified, two more differentiated clusters: **E**arly **D**igit **C**ondensations (**EDC**: *Hoxd13+; Sox9+)* and **L**ate **D**igit **C**ondensations (**LDC**: *Irx1+*). Finally, in-between proximal and distal regions (*Shox2+* and *Hoxd13+*), we found a cluster of condensating cells that we considered to be the **M**e**s**opodium (**Ms**: *Sox9+*; *FoxcI+, Gdf5+*) and that thus corresponds to ankles or wrists (**Fig. 2C-D, Supplementary Table S1**). To better understand the links between the different clusters, we ran an RNA velocity analysis in the hindlimb dataset (**Fig. 2E**) (La Manno et al. 2018; Bergen et al. 2020). We found that in the proximal limb a set of *Irx5*-expressing cells located within the PPP and ICT clusters are progenitor for the more differentiated proximal clusters such as TP and PC (**Fig2D, 2E**) (Li et al. 2014). In the distal limb, DP and DPP clusters appear to be progenitor for EDC and then LDC. The Mesopodium cluster originates from both proximal (PPP-ICT) and distal (DP-DPP) progenitor clusters, confirming its proximo-distal origin.

**Figure 2.**
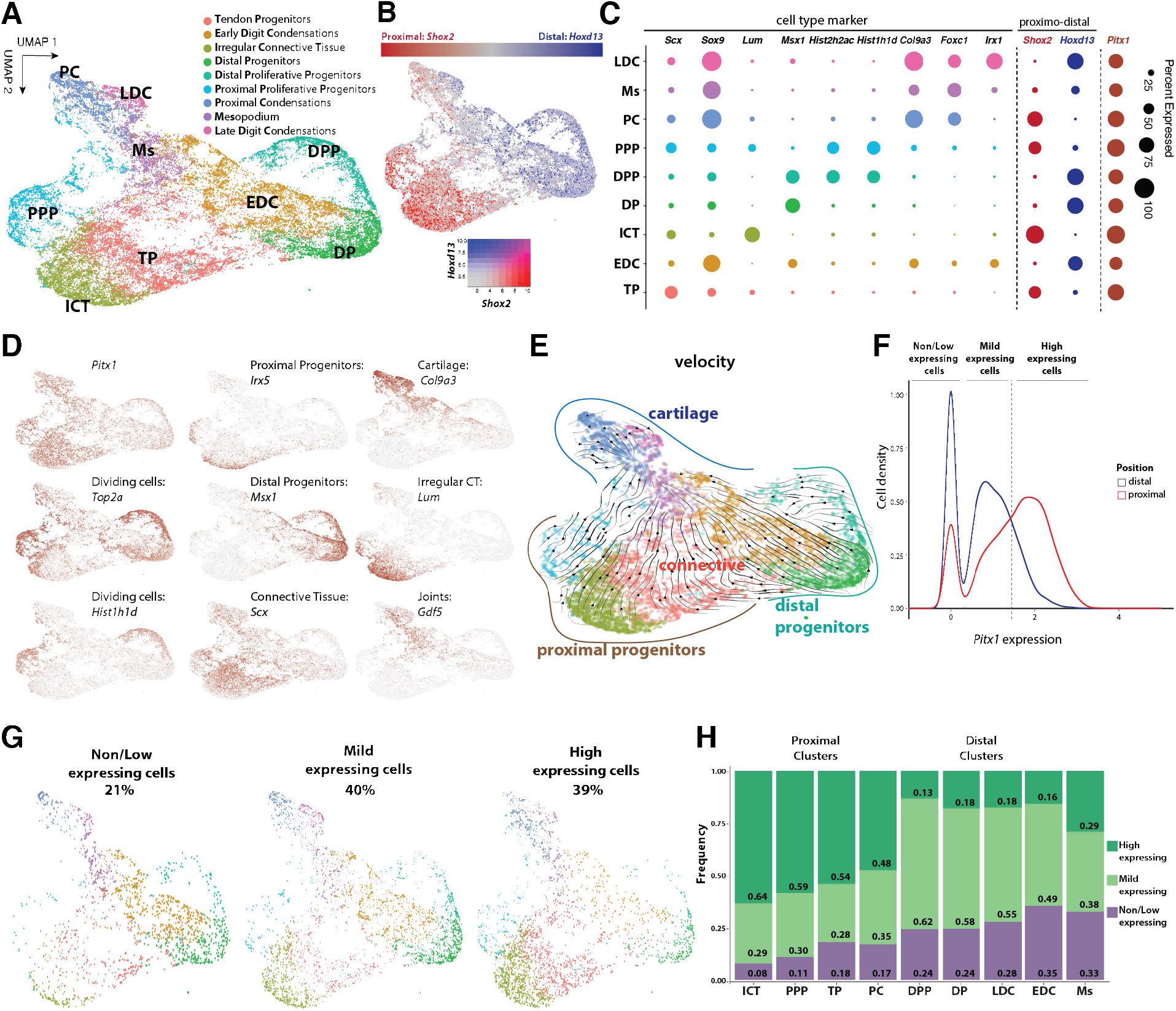
*Pitx1* expression in hindlimbs: **A**. UMAP of the re-clustering of mesenchymal cells from all datasets. **B**. Distribution of *Shox2* (proximal) and *Hoxd13* (distal) marker. **C**. Representative marker genes for each cluster. The dot size corresponds to the percentage of cells that express a given marker in the hindlimb wildtype dataset. **D**. Expression distribution of selected marker genes across the UMAP **E**. RNA-velocity analysis of hindlimb wildtype mesenchymal clusters. Note that the different differentiated cell cluster (upper part) derive from proximal and distal progenitors’ clusters (bottom part). **F**. *Pitx1* expression density plot in the proximal (red line) and distal clusters (green line) in the hindlimb wildtype dataset. Definition of the three types of *Pitx1*-expressing cells: non/low-(<0.3 *Pitx1* expression levels), mild-(0.3-1.4), high-expressing (>1.4). **G**. Hindlimb wildtype cells distribution across the clusters in the UMAP space based on their *Pitx1* levels of transcription. **H**. Hindlimb wildtype cells proportions according to *Pitx1* expression level across mesenchymal clusters.

We then asked whether *Pitx1* is differentially expressed among clusters in hindlimb wildtype. Overall, we found *Pitx1* expressed in all mesenchymal clusters, yet with a proximal preference (**Fig. 2D, 2F-G**). We then classified *Pitx1* expressing cells in three categories: non/low-expressing (21 % of the hindlimb wildtype cells), mild-expressing (40 % of cells), and high-expressing (39 % of cells) (**Fig. 2F-G)**. Expectedly, we found that a majority of high expressing cells are located in proximal clusters (PPP, TP, ICT, PC) and a majority of mild-expressing cells in distal clusters (DP, DPP, EDC, LDC) (**Fig. 2F-H)**. We also observed that the Ms cluster, previously identified as a cluster originating from the proximal and distal cell-types, is formed by a similar distribution of high-expressing (proximal) and mild-expressing (distal) cells in line with a proximo-distal origin (**Fig 2H**).

### *Pitx1* expression levels associate with global change in regulatory landscape acetylation

Next, we explored how cells can achieve distinct *Pitx1* transcriptional outputs. Practically, we asked whether high- and mild-expressing cells use a distinct *Pitx1* enhancer repertoire to account for the different expression levels. We sorted the two cell populations from *Pitx1*^*GFP*^ hindlimbs by GFP intensities: GFP+-(mild-expressing) and GFP++ (high-expressing) and performed H3K27ac ChIP-seq on the two positive populations (**Fig. 3A and Fig. S3**). As expected from the single-cell analysis, high-expressing GFP++ cells were mostly derived from proximal limbs as demonstrated by the anterior activity of the *HoxA* and *HoxD* clusters and by *Shox2* activity (**Fig. 2G, 3B, 3C and Fig. S3**). In contrast, mild-expressing cells GFP+-where enriched for distal cell markers such as *Hoxa13, Evx1, Msx1 and Hoxd13 and Evx2* (**Fig. 3C, Fig. S3**).

**Figure 3.**
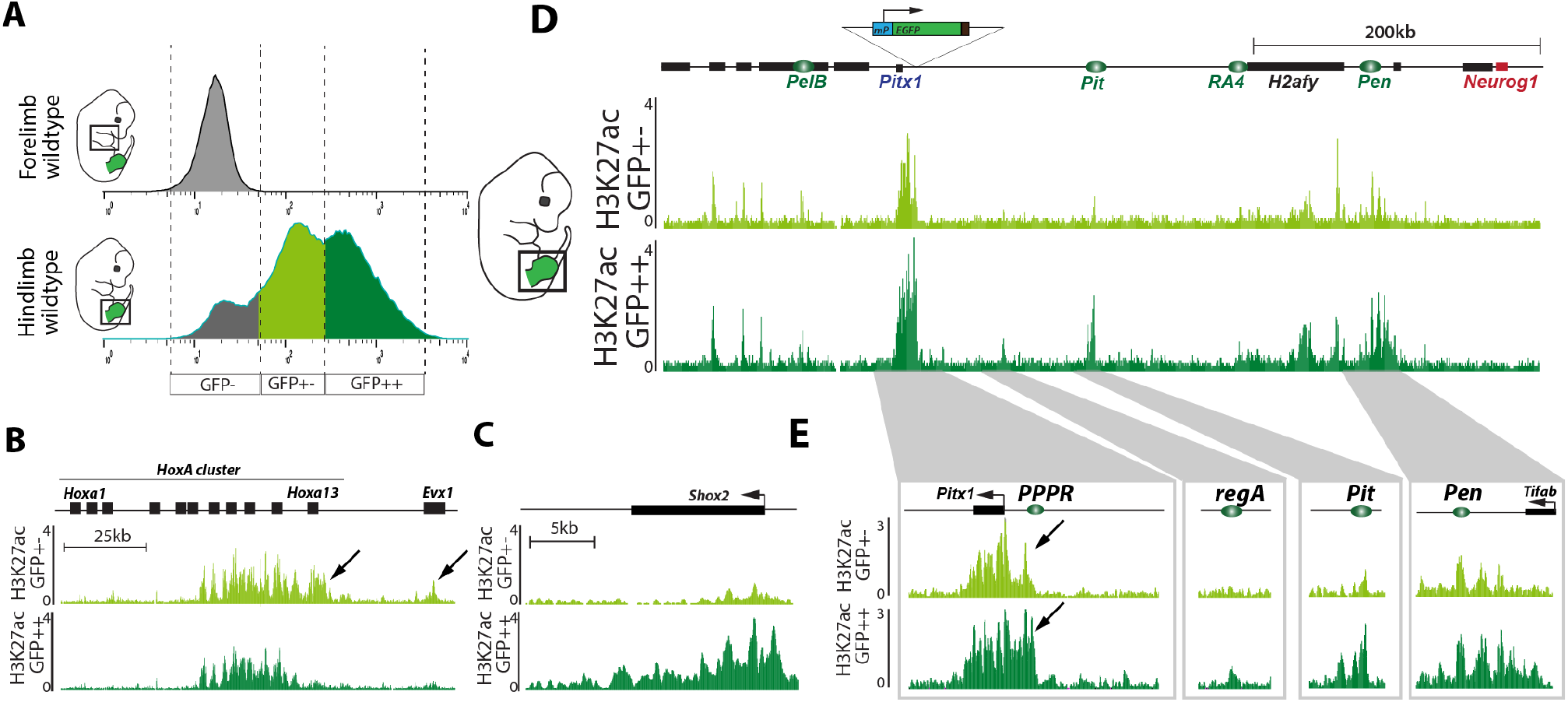
High- and mild-expressing *Pitx1* regulatory landscape activities: **A**. FACS sorting of wildtype *Pitx1*^*GFPs/GFPs*^ forelimb and hindlimbs. Note the apparent EGFP high-(dark green) and mild-expressing (light green) populations. **B**. H3K27ac ChIP-seq in mild- and high-expressing hindlimb cell populations at the *HoxA* cluster. Note the distal *Hoxa13* and *Evx1* genes activities (black arrows) in the mild active cells. **C**. H3K27ac ChIP-seq at the *Shox2* locus. Note the activity in the highly active cells. **D**. H3K27ac ChIP-seq at the *Pitx1* locus. Note that enhancers are active in both mild and high expressing cells, yet with a few regions marked only in high-expressing cells. **E**. H3K27ac profile on *Pitx1* gene body, *Pitx1* Proximal Promoter Region (PPPR, see black arrow), region A (regA), *Pit* and the *Pen* enhancer.

In both mild- and high-expressing cells, the previously characterized *Pitx1* enhancer repertoire - *PelB, Pit, RA4* and *Pen* - was found marked by H3K27ac. Yet, in high-expressing cells, stronger H3K27ac signal was found at these elements concomitantly with a drastic increase at two specific regions: the *Pitx1* proximal promoter region (*PPPR*) and the region A (*regA*) (**Fig. 3D and 3E**) (Kragesteen et al. 2018). This data shows that *Pitx1* regional expression differences across hindlimbs associate with a progressive increase of its *cis*-regulatory landscape activity rather than different repertoires of enhancers. This results further re-enforce the idea that the fundamental unit of *Pitx1* regulation is the landscape as a whole rather than individual enhancers.

### *Pen* deletion increases *Pitx1* non/low-expressing cells and alters limb cell composition

Seeing the coordination between regulatory units at the locus to modulate gene expression we sought to test how the deletion of one of them influences the overall unity of the locus. Therefore, we took advantage of the *Pitx1* EGFP sensor and scRNA-seq to track how the homozygous deletion of the *Pen* enhancer affects the hindlimb *Pitx1* locus activity. Using scRNA-seq, we found that the *Pen* deletion induces a significant 29% loss of *Pitx1* expression (adjusted p-value=1.75e-96(Wilcoxon Rank Sum test)) featured by a decrease in *Pitx1* high-expressing cells and a strong increase in low/non-expressing cells (**Fig. 4A**). Across hindlimb mesenchymal cells, the proportion of non/low-expressing cells was indeed raised from 21% in wildtype to 35% in *Pitx1*^*Pen-/Pen-*^. We observed a similar effect when we quantified EGFP fluorescence of *Pitx1*^*GFP;ΔPen*^ mutant hindlimbs where high-fluorescent cells are lost and low/non-fluorescent cells are increased (**Fig. 4B-C**). In this case, the proportion of GFP-cells raised from 8% in *Pitx1*^*GFP*^ to 16% in *Pitx1*^*GFP;ΔPen*^ at E12.5 and from 12% to 29% at E13.5 (**Fig. S4A-B**). Yet, from the fluorescence distribution, it appears that a higher average EGFP level is visible within the GFP-gating, thereby indicating the presence of a higher proportion of low-expressing cells (**Fig. S4A-B)**. In summary, the two approaches show that behind the weak average loss of *Pitx1* expression, a strong increase of non/low-expressing cells in mutant hindlimbs could account for the clubfoot phenotype seen in these animals (Kragesteen et al. 2018).

**Figure 4.**
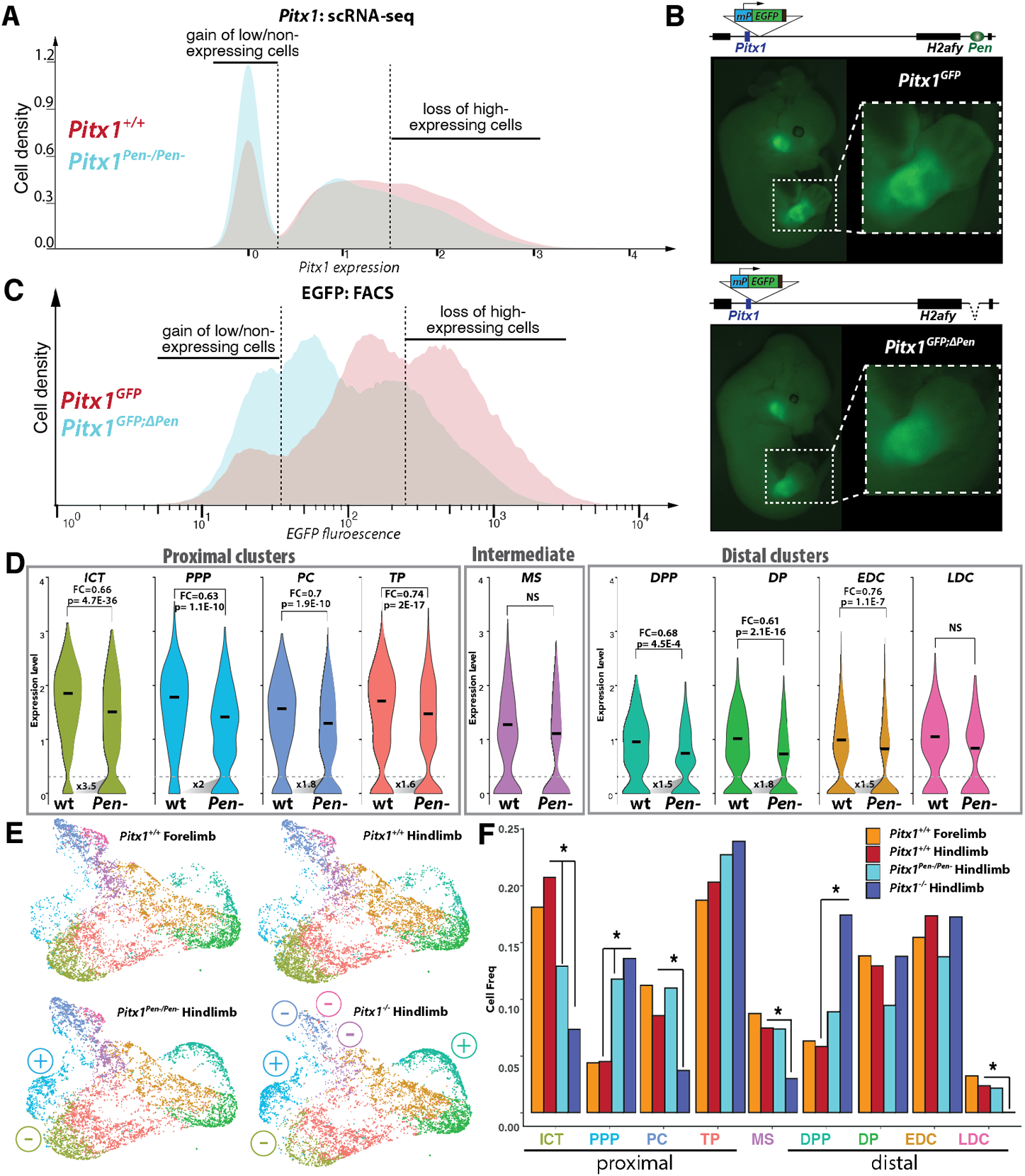
Influence of the Pen deletion on Pitx1 expression in hindlimb cell population. **A**. *Pitx1* expression distribution across wildtype (red) and *Pitx1*^*Pen-/Pen-*^(cyan) hindlimb cells shows an increased proportion of non/low-expressing mutant cells and a decrease proportion of high-expressing cells. **B**. EGFP expression pattern in *Pitx1*^*GFP*^ and *Pitx1*^*GFP;ΔPen*^ in E12.5 embryos. **C**. FACS profile of *Pitx1*^*GFP*^ (red) and *Pitx1*^*GFP;ΔPen*^ (cyan) hindlimbs shows an increased number of EGFP non/low-expressing cells as well as a decrease of EGFP high-expressing cells. **D**. *Pitx1* expression across all clusters in *Pitx1*^*GFP*^ and *Pitx1*^*GFP;ΔPen*^ hindlimb. At the base of the distribution, the fold change in non/low-expressing cell number between wildtype and mutant is shown. Note the strong loss of expression and the accumulation of non/low-expressing cells in ICT and PPP clusters. **E.F**. UMAP **(E)** and quantification **(F)** of mesenchyme cell type proportions across conditions. (+) and (-) symbols indicate increase or decrease in cell proportions, stars indicate p<0.05.

We further asked if this alteration in expression was equally distributed among various hindlimb cell-types or if some populations were more specifically affected. All clusters with the exception of the Ms and the LDC showed a significant loss of *Pitx1* expression ranging from 24% to 39% (**Fig. 4D)**. With respect to the proportion of non/low-expressing cells, we saw that proximal clusters showed a preferential 2.1-fold enrichment of non/low-expressing cells (13% to 28%) in comparison with distal cells (1.6-fold, 29% to 45%) (**Fig. S5A-B)**. We then computed the increase of non/low-expressing *Pitx1* cells in each cluster and saw that two proximal clusters in particular, ICT and the PPP, showed a 3.5- and 2-fold increase in *Pitx1* non/low-expressing cells, respectively. It is important to note that in both clusters the vast majority of cells usually express *Pitx1* at a high level (**Fig. 2H, S6**). Other clusters showed 1.5-to 1.8-fold increase in *Pitx1* non/low-expressing cells. In conclusion, we found that proximal, high-expressing clusters are more affected by the enhancer deletion than distal, mild-expressing clusters. We then investigated if this differential alteration of *Pitx1* expression among hindlimb cell population affected the proportion of cells within the clusters.

As a positive control for the effect of *Pitx1* loss-of-function in *Pitx1*^*Pen-/Pen-*^ embryos, we took advantage of two datasets that do not express *Pitx1* at all: wildtype forelimbs and *Pitx1*^*-/-*^ hindlimbs. First, by comparing wildtype fore- and hindlimbs, we did not observe any significant change in the proportion of cell-types in either tissue, suggesting that the developmental origin of their cell populations is identical despite the obvious structural differences between arms and legs (**Fig. 4E, 4F**). In contrast, *Pitx1*^*-/-*^ hindlimbs display a heterochronic phenotype, featuring an increase in progenitor cells in both the proximal and distal regions of the limb (PPP and DPP) while a concurrent decrease is seen in several differentiated cell types in proximal and distal limbs (ICT, PC, Ms and LDC) (**Fig. 4E, 4F**). Remarkably, the loss of the *Pen* enhancer resulted in a similar effect but only in the proximal limb cell clusters (**Fig. 4E and 4F**). Specifically, the proportion of PPP cells increased in *Pitx1*^*Pen-/Pen-*^ hindlimbs as the proportion of ICT cells decreased. This strong effect shows that the loss of *Pitx1* in these clusters is enough to perturbate the proportion of cells that compose them. Moreover, this data suggests that a failure to reach an appropriate cell-type specific gene expression level is at the basis of the clubfoot phenotype.

As *Pitx1* has been shown to have both indirect and direct downstream effects, we further investigated if we could detect such changes in *Pitx1*^*Pen-/Pen-*^ hindlimbs. In particular, it has been shown that *Tbx4* mediates the *Pitx1*-effect on hindlimb buds growth rate (Duboc and Logan 2011). As anticipated, we found a clear downregulation of the *Tbx4* in all clusters with except of PC, Ms and LDC in both *Pitx1*^*- /-*^ and *Pitx1*^*Pen-/Pen-*^ hindlimbs (**Fig. S7A-C**). Moreover, in *Pitx1*^*Pen-/Pen-*^hindlimbs, the magnitude of *Tbx4* loss-of-expression followed the one of *Pitx1*. In particular, we saw a strong decrease of *Tbx4* expression and an increase of *Tbx4* non-expressing cells in ICT and PPP clusters. This suggests that in the case of the *Pen* deletion an important fraction of the pathological effect might be conveyed *via Tbx4*.

### The *Pen* enhancer contributes to *Pitx1* regulatory landscape activation

The establishment of the active *Pitx1* chromatin landscape includes changes in 3D conformation and the acetylation of specific *cis*-regulatory elements. Therefore, we asked whether the *Pen* enhancer itself is required to establish these features and specifically if its deletion would impact them.

In sorted GFP+ and GFP-in *Pitx1*^*GFP;ΔPen*^, we first assessed using RNA-seq whether we could observe similar changes in cellular identity upon *Pen* enhancer loss as the one previously described with scRNA-seq. As expected, we could observe in GFP-cells the accumulation of mesenchymal markers (*Prrx1, Twist1*) with a particular enrichment for ICT markers (*Col3a1, Col1a1, Col1a2, Lum)* (**Fig. 5A, Supplementary Table S3)**. Because of the accumulation of cells losing *Pitx1* activity, and the consequent dilution of non-mesenchymal clusters, we also observed a decrease in satellite marker genes including epithelium (*Wnt6, Krt14*) and muscle (*Ttn*) markers. In GFP+ cells, we did not observed a clear change in identity markers suggesting that the cell type composition is similar between wildtype and mutant high-expressing cells (**Supplementary Table S4**). We then tested whether these high-expressing *escaping* cells display an adaptive mechanism to accommodate the *Pen* enhancer loss.

**Figure 5.**
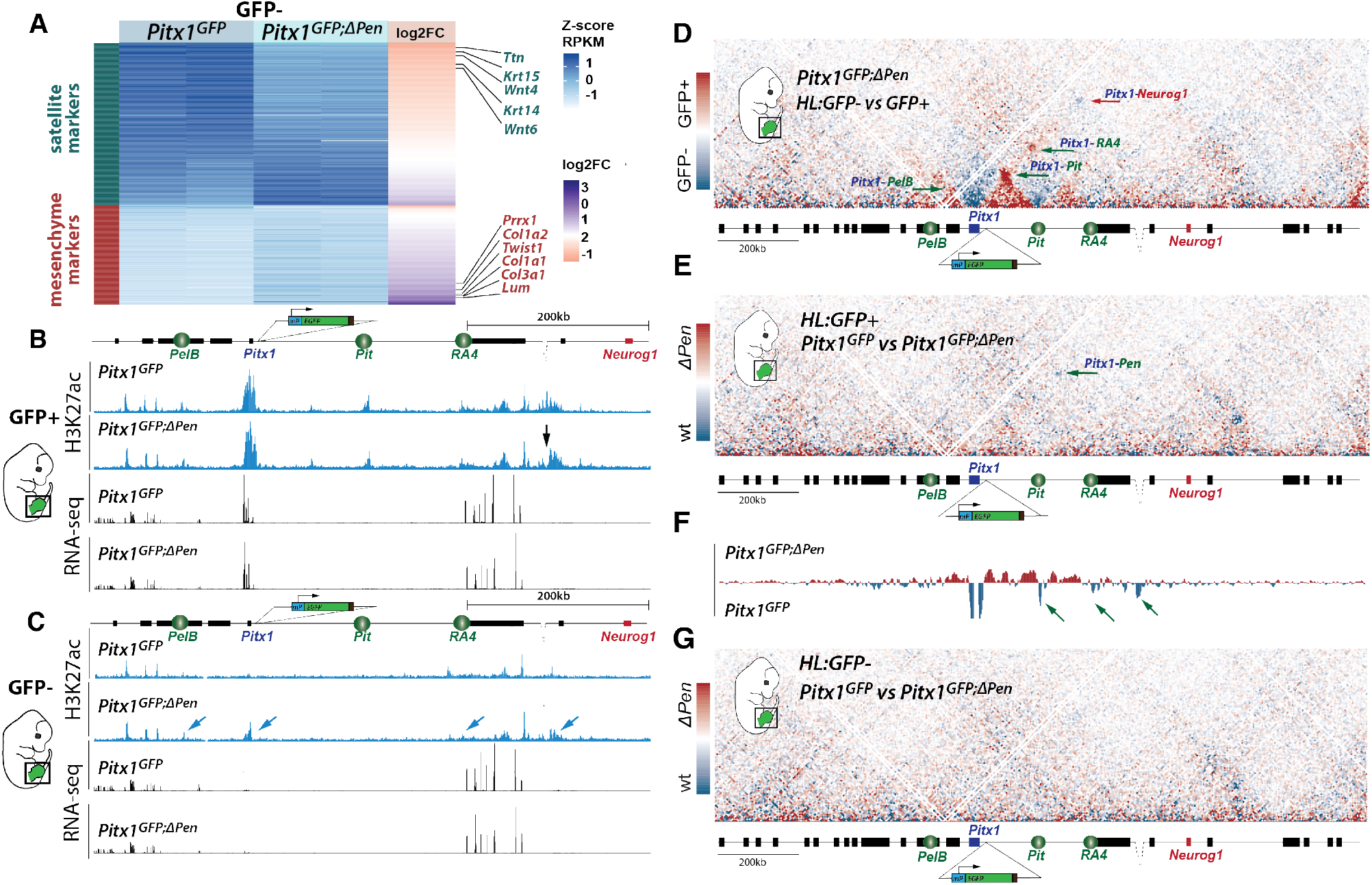
Single enhancer deletion results in inefficient regulatory landscape activation. **A**. Log2 fold change and RPKM of mesenchymal (orange) and satellite (darkgreen) marker genes in *Pitx1*^*GFP*^ and *Pitx1*^*GFP;ΔPen*^ GFP-hindlimbs cells. Note the decrease in satellite markers and the increase in mesenchymal markers in *Pitx1*^*GFP;ΔPen*^ GFP-cells **B**. H3K27ac ChIP-seq and RNA-seq at the *Pitx1* locus in GFP+ hindlimb cells in *Pitx1*^*GFP;ΔPen*^ background. Note the loss of the *Pen* enhancer region (black arrow). **C**. H3K27ac ChIP-seq and RNA-seq at the *Pitx1* locus in GFP-hindlimb cells in *Pitx1*^*GFP;ΔPen*^ background. Note the acetylation of *Pitx1* promoter and enhancers cells (blue arrows) and the weak Pitx1 transcription. **D**. C-HiC in subtraction between GFP-and GFP+ sorted *Pitx1*^*GFP;ΔPen*^ hindlimb (HL) cells. **E**. C-HiC in subtraction between *Pitx1*^*GFP*^ GFP+ hindlimb cells and *Pitx1*^*GFP;ΔPen*^ GFP+ cells. **F**. Subtraction track of virtual 4C between *Pitx1*^*GFP*^ and *Pitx1*^*GFP;ΔPen*^ GFP+ hindlimb cells from the *Pitx1* viewpoint. Note the loss of interaction between *Pitx1* and its telomeric enhancers (*Pit, RA4* and *Pen*). **G**. C-HiC in subtraction between *Pitx1*^*GFP*^ GFP-hindlimb cells and *Pitx1*^*GFP;ΔPen*^ GFP-cells.

Specifically, we performed H3K27ac ChIP-seq in the escaping GFP+ cells and in the increased GFP-cell population. In GFP+ cells, we observed a distribution of H3K27ac over the landscape that was virtually identical to wildtype GFP+ hindlimbs cells, with the exception of the *Pen* enhancer itself (**Fig. 1E, 5B**). This result suggests that the *Pitx1* expressing cells in the *Pen* deletion background use the same enhancer repertoire as the wildtype expressing cells and thus, do not use an alternative regulatory landscape. Moreover, we observed the same average *Pitx1* expression level in wildtype and mutant GFP+ cells (**Supplementary Table S4**). In GFP-cells deleted for *Pen*, in contrast to wildtype cells, we observed ectopic acetylation of the *Pitx1* promoter as well as of the *RA4* and *PelB* enhancers (**Fig. 5C)**. These activities are likely caused by the relocation in the GFP-gating of cells that would normally express *Pitx1* but failed to establish a fully active landscape in the absence of *Pen*. As expected from the marginal increase in EGFP fluorescent cells previously described (**Fig. 4C, S4**), we also observed a marginal but significant increase in *Pitx1* expression (FC=1.6, padj=0.0026) far from the expression level observed in transcriptionally active cells **(Fig. 5C, Supplementary Table S3)**.

We then measured how the lack of *Pen* affects the locus 3D structure dynamics in *Pitx1*^*GFP;ΔPen*^ hindlimbs. First, GFP+ and GFP-*Pitx1*^*GFP;ΔPen*^ hindlimb cells displayed differences similar to their wildtype *Pitx1*^*GFP*^ active and inactive counterparts. This suggests that escaping high-expressing hindlimb *Pitx1*^*GFP;ΔPen*^ cells do not require *Pen* to establish an active 3D conformation (**Fig. 1F, 5D, S8A**). We thus asked whether these cells bare an alternative chromatin structure than wildtype ones to compensate for the loss of *Pen*. By comparing wildtype and *Pen*-deleted GFP+ cells we saw no major differences (**Fig. 5E, S8B**). Yet, using virtual 4C, we saw a slight reduction of contact between the *Pitx1* promoter and *Pit/RA4* in GFP+ cells (**Fig. 5F**). This suggests that the remaining high-expressing cells do not necessarily undergo a strong adaptive structural response to the loss of *Pen* to ensure high *Pitx1* expression. Finally, we asked whether the relocated *Pitx1*^*GFP;ΔPen*^ GFP-non-expressing cells, that bear ectopic promoter and enhancer acetylation, also display features of an active 3D structure (**Fig. 5G, S8C**). However, we did not observe any change in the *Pitx1* locus conformation in these cells in comparison to wildtype non-expressing cells. This shows that despite bearing some regulatory activity, the locus is unable to undertake its active 3D structure and therefore to efficiently transcribe *Pitx1*. In conclusion, the *Pen* enhancer is necessary to ensure that all the cells with active enhancers at the *Pitx1* locus undergo a robust transition toward a structurally and transcriptionally active landscape (**Fig. 6**).

**Figure 6.**
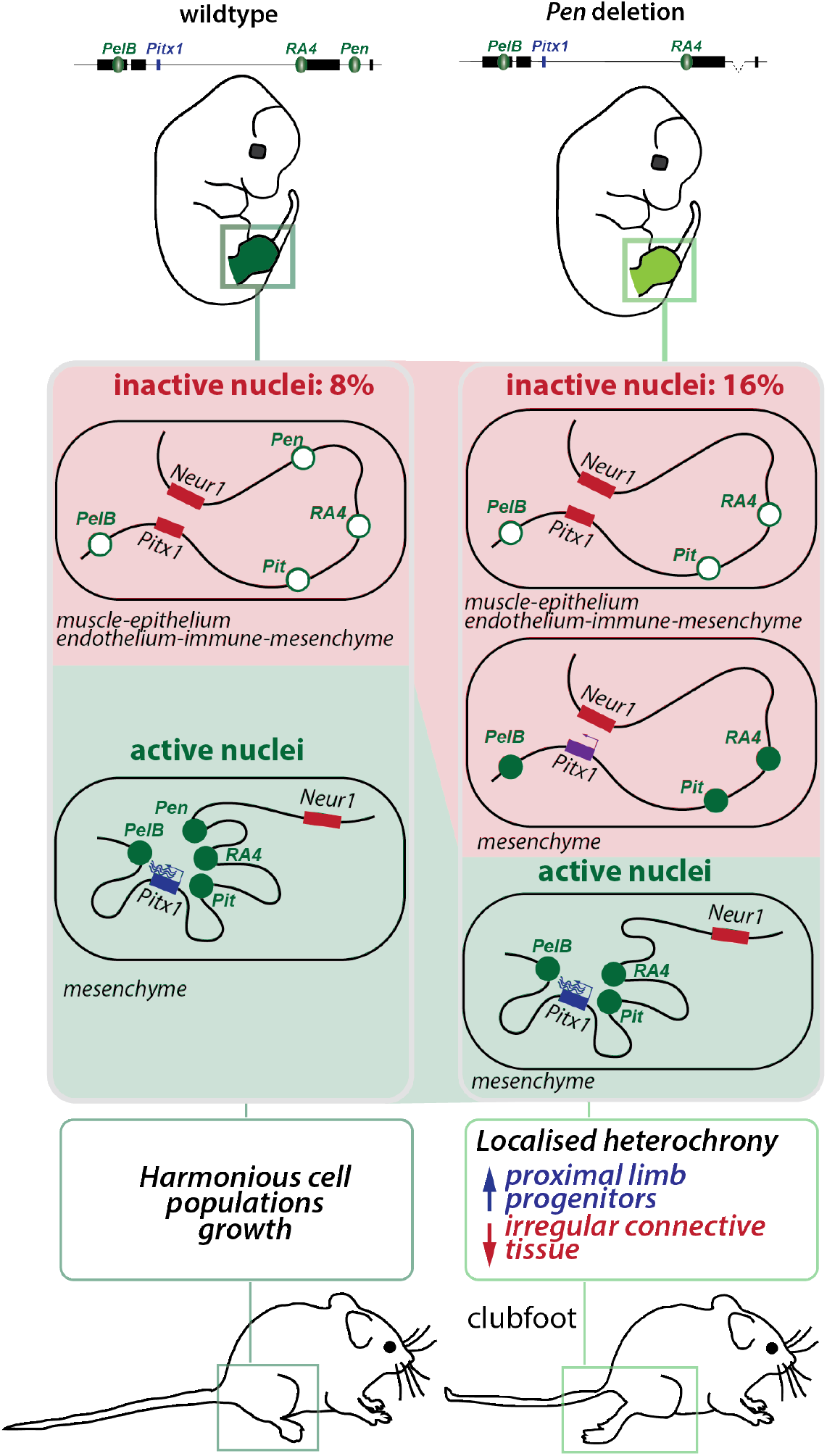
Model. In wildtype hindlimb tissues (left panel) 8% of the nuclei, mostly in non-mesenchymal cell types, display a completely repressed locus and 3D chromatin structure. In active nuclei, the situation is inverted with an active regulatory landscape, an active 3D chromatin structure and strong *Pitx1* transcription. In contrast, in hindlimb lacking the *Pen* enhancer (right panel), 16% of the cells are lacking *Pitx1* transcription. Among these cells, some display a partially active regulatory landscape. These cells, that have failed to establish an active 3D structure and a strong *Pitx1* transcription, are of mesenchymal origins with a preference for ICT. The remaining active cells in mutant hindlimbs appear to display wildtype expression levels despite lacking an important enhancer. Generally, the effect of the enhancer deletion on the limb outgrowth is a disharmonious proportion of cell type including a gain of PPP and a decrease of ICT, leading to a clubfoot phenotype.

## Discussion

In this work we have shown that hindlimb cells display several states of *Pitx1* regulatory activities. In active cells, all enhancers are marked with the active H3K27ac chromatin modification and are contacting the *Pitx1* promoter. In contrast, in inactive cells, we could not observe partial regulatory activities, i.e. neither enhancer acetylation nor enhancer-promoter interactions. This shows that the locus follows a bimodal behavior where the regulatory landscape as a whole acts on *Pitx1* transcription. Indeed, a common set of coordinated enhancers are active in both proximal *Pitx1* high-expressing and distal *Pitx1* low-expressing cells. In fact, the *Pitx1* regulatory landscape acts here similarly to what was previously define as a holo-enhancer, where the whole region seems to work as a *coherent regulatory ensemble* (Marinic et al. 2013). In this perspective, *Pitx1* expression levels are adjusted by the entire landscape. This is what we observed in high *Pitx1*-expressing proximal cells where the same enhancer set than in distal cells displays a higher enrichment for active H3K27ac chromatin mark along with a few proximal-specific regions that are more enriched for H3K27ac.

Here we have tested how the loss of one the regulatory element, the *Pen* enhancer, affects the establishment of the active landscape. As some *escaping* cells can induce *Pitx1* regulatory landscape *activation* without *Pen*, many very low to non *Pitx1*-expressing cells accumulate in hindlimbs. The latter cells bear high enrichment of H3K27ac at the *Pitx1* promoter and at several of its enhancers (**Fig. 6**). However, despite the presence of this active modification, the *Pitx1* locus does not adopt an active 3D chromatin folding but maintains the hallmarks of its inactive configuration. In fact, these accumulated low/non-expressing cells are seemingly stuck in a *limbo* between activity and repression and show the importance of the coordinated action of enhancer activity and 3D chromatin changes to achieve sufficient transcriptional strength. Therefore, we hypothesize that the role of *Pen* is not to act as a pattern-defining enhancer but rather as a support enhancer that ensures a robust transition of cells towards a fully active landscape and therefore a strong *Pitx1* transcription. In fact, *Pen* is a good model to understand the fundamental role of many enhancers that were characterized with a diverging activity than the gene they control (Visel et al. 2007; Ruf et al. 2011; Symmons and Spitz 2013). This “class” of enhancers would therefore govern the cooperativity of the regulatory landscape of their respective locus without defining by themselves its expression specificities.

Changes in the number of cells that express *Pitx1* in the hindlimb have strong phenotypical consequences. In fact, the complete loss of *Pitx1* induces a strong increase in proximal and distal progenitor cells concomitantly with a loss of differentiated cell types, overall altering the proportion of specific cell clusters in hindlimbs. The global increase in progenitors indicates a heterochrony in limb development that ultimately results in a reduction of the limb size and the loss of some limb structures such as the patella. In the case of the *Pen* enhancer deletion, we saw a dramatic enrichment of *Pitx1* low/non-expressing cells in PPP and ICT clusters, two clusters where cell numbers were proportionally altered. This cellular proportion alteration might be mediated *via Tbx4* which is lost in these clusters and was shown to be important for limb outgrowth (Duboc and Logan 2011). Here, the particularly strong effect of the *Pen* deletion on the ICT cell proportion pinpoints these cells as the origin of the clubfoot phenotype seen in mice lacking the enhancer. In fact, ICT that comprises muscle connective tissue, has been reported as a major driver of limb morphogenesis and our data suggest that it might be at the base of the clubfoot phenotype (Besse et al. 2020; Helmbacher and Stricker 2020). Finally, despite lacking *Pitx1* expression as well, forelimb cell clusters are present in the same proportion as hindlimb ones. This suggests that the role of *Pitx1* in hindlimb is mirrored by other genes in forelimbs, such as *Tbx5*, that account for a normal harmonious outgrowth of the various cell populations. Indeed, *Tbx5* loss of expression in the ICT population alters muscle and tendons patterning causing the mice to hold the paw in a supine position, leading them to walk on the edge or dorsal surface of the paw, resembling a clubfoot phenotype (Besse et al. 2020).

Our characterization of a single enhancer loss-of-function mutant at a cell subpopulation levels opens the way to study the effect of other regulatory mutations with the same resolution and, in particular, of gain-of-function mutations. Such approaches will enable to select particular cell-subpopulations that show ectopic expression in comparison to neighboring cells that bear the same mutation but no ectopic expression. This will facilitate a precise definition of features that are permissive for transcriptional gain-of-function and will be an important tool to further investigate the relationship between 3D structure, chromatin modifications and gene transcriptional activation.

## METHODS

### CELL CULTURE AND MICE

#### CRISPR/Cas9 Engineered Alleles

Genetically engineered alleles were generated using the CRISPR/Cas9 editing according to (Kraft et al. 2015). Briefly, sgRNAs were designed using the online software Benchling and were chosen based on predicted off-target and on-target scores. All sgRNAs and target genomic locations for CRISPR-Cas9 can be found in Supplementary Table S5. SgRNAs were then sub-cloned in the pX459 plasmid from Addgene and 8 μg of each vectors was used for mESCs transfection. mESCs culture and genetic editing followed standard procedure (Andrey and Spielmann 2017).To construct the *Pitx1*^*GFP*^ mESCs clone, a the lacZ sensor from (Kragesteen et al. 2018) was adapted by exchanging the LacZ by an EGFP cassette. The sgRNA was designed to target CRISPR-Cas9 to chr13:55935371-55935390 (Supplementary Table S5). Cells were transfected with 4μg of EGFP-cassette and 8μg of pX459 vector containing the sgRNA.

#### Aggregation of mESC

Embryos were generated by tetraploid complementation from G4 ESCs (George et al. 2007; Artus and Hadjantonakis 2011). Desired mESCs were thawed, seeded on CD1 feeders and grown for 2 days before the aggregation procedure. Donor tetraploid morula are from B6D2F1 background and embryos were transferred into foster CD1 female.

#### Animal Procedures

All animal procedures were in accordance with institutional, state, and government regulations (Canton de Genève authorisation: GE/89/19).

#### Single-Cell RNA-seq dissociation

Two replicates of fore and hindlimb buds of E12.5 wildtype embryos and hindlimb buds of mutant embryos (*Pitx1*^*Pen-/Pen-*^, *Pitx1*^*-/-*^) were micro-dissected and incubated for 12 minutes in 400μl trypsin-EDTA 0.25% (Thermo Fischer Scientific, 25300062), supplemented with 40μl of 5% BSA. During incubation tissues were disrupted by pipetting after 6 minutes of incubation and at the end of the 12 minutes. Trypsin was then inactivated by adding 2x volume of 5% BSA and single cell suspension was obtained by passing cells in a 40μm cell strainer. Cells were then spun at 250g for 5 minutes at 4° and resuspended in 1%BSA in PBS. Cells were then counted using an automatized cell counter and a 1% BSA 700cells/ul suspension was prepared. 10ul of this solution was used as input for the 10X Genomics library preparation.

#### Single-Cell library preparation

Single-cell libraries were prepared using the Chromium Single Cell 3’ GEM, Library & Gel Bead Kit v3 following the manufacture’s protocol (10X Genomics, PN-1000075). Briefly, Gel beads in EMulsion (GEMs) are generated by combining Single Cell 3′ v3 Gel Beads, a Master Mix containing cells, and Partitioning Oil onto Chromium Chip B. Incubation of the GEMs produced from the poly-adenylated mRNA barcoded, full-length cDNA. Immediately, gel beads are dissolved and cDNA is amplified via PCR followed by library construction and sequencing. Libraries were paired-end sequenced on a HiSeq 4000. On average, 7000 cells were loaded on the Chromium Chip and between 25000-35000 mean reads were obtained.

#### Whole-mount in situ hybridization (WISH)

*Pitx1* WISH were performed on 40-45 somite stage mouse embryos (E12.5) using a digoxigenin-labeled *Pitx1* antisense riboprobe transcribed from a cloned *Pitx1* probe (PCR DIG Probe Synthesis Kit, Roche), as previously described in (Kragesteen et al. 2018).

#### Tissue collection and cell preparation for FACS-sorting

Forelimb and hindlimb buds from embryos with 40-45 somites (E12.5) were dissected in cold PBS solution. After PBS removal, a single cell suspension was achieved by incubating the limb buds in 400uL Trypsin-EDTA (Thermo Fischer Scientific, 25300062) for 12’ at 37°C in a Thermomixer with a resuspension step at the 6’ mark. After blocking with one volume of 5% BSA (Sigma Aldrich, A7906-100G), cells were passed through a 40μm cell strainer for further tissue disruption and another volume of 5% BSA was added to the cell strainer to pass leftover cells. Cells were then centrifuged at 400g for 5’ at 4°C and, after discarding the supernatant, they were resuspended in 1% BSA for cell sorting. 5mM of NaButyrate were added to the BSA when planning for subsequent fixation for H3K27Ac-ChIP.

#### Cell sorting

Cell populations were isolated using fluorescent-activated cell sorting (FACS) using the Beckman Coulter MoFlo Astrios with GFP laser (excitation wavelength 488nm). After removal of dead cells with Draq7 dye and removal of doublets, following standard protocol, cells were gated for sorting as can be seen in **FigS1A**. Flow cytometry analysis to obtained GFP histograms was performed with the FlowJo^™^ Software (version 10.6.1).

#### Cell processing for RNA-seq, ChIP-seq and Capture-HiC

##### ChIP and Capture-HiC

After sorting, cells were centrifuged for 5’ at 400g at 4°C and supernatant was discarded. Cells for ChIP-seq and Capture-HiC were resuspended in 10% FCS/PBS and fixed in 1% formaldehyde for ChIP and 2% for Capture-HiC at room temperature. The fixation was blocked by the addition of 1.25M glycine, cells were isolated by centrifugation (1000g, at 4°C for 8’), resuspended in cold lysis buffer (10 mM Tris, pH 7.5, 10 mM NaCl, 5 mM MgCl2, 0.1 mM EGTA, Protease Inhibitor (Roche, 04693159001)) and incubated on ice for 10’ to isolate the cell nuclei. The nuclei were isolated by centrifugation (1000g, at 4°C for 3’), washed in cold 1X PBS, centrifuged again (1000g, at 4°C for 1’) and stored frozen at −80°C after removal of the PBS supernatant.

##### RNA-seq and library preparation

After sorting, cells were centrifuged for 5’ at 400g at 4°C, supernatant was discarded and cells frozen at −80°C. Total RNA from 1,5 x10^5^ cells was isolated using the RNeasy Micro Kit (QIAGEN, ID:74004) following manufacturer’s instructions and then stored frozen at −80°C. Total RNA was quantified with a Qubit (fluorimeter from Life Technologies) and RNA integrity assessed with a Bioanalyzer (Agilent Technologies). The SMART-Seq v4 kit from Clontech was used for the reverse transcription and cDNA amplification according to manufacturer’s specifications, starting with 5 ng of total RNA as input. 200 pg of cDNA were used for library preparation using the Nextera XT kit from Illumina. Library molarity and quality was assessed with the Qubit and Tapestation using a DNA High sensitivity chip (Agilent Technologies). Libraries were pooled at 2 nM and loaded for clustering on a Single-read Illumina Flow cell for an average of 35 mio reads / library. Reads of 50 bases were generated using the TruSeq SBS chemistry on an Illumina HiSeq 4000 sequencer.

##### ChIP-seq and library preparation

5×10^5^ fixed nuclei were sonicated to a 200-500bp length with the Bioruptor Pico sonicator (Diagenode). H3K27Ac ChIP was (Diagenode C15410174) was performed as previously described (Lee et al. 2006; Paliou et al. 2019) with the addition of 5mM of Na-Butyrate to all buffers. Libraries were then prepared following the Illumina ChIP TruSeq protocol and sequenced as 50bp single-end reads on a illumina HiSeq 4000. Libraries were prepared starting with below <10ng quantities of ChIP-enriched DNA as starting material and processed with the Illumina TruSeq ChIP kit according to manufacturer specifications. Libraries were validated on a Tapestation 2200 (Agilent) and a Qubit fluorimeter (Invitrogen – Thermofisher Scientific). Libraries were pooled at 2 nM and loaded for clustering on a Single-read Illumina Flow cell. Reads of 50 bases were generated using the TruSeq SBS chemistry on an Illumina HiSeq 4000 sequencer.

##### Capture-HiC and library preparation

3C libraries were prepared as previously described (Paliou et al. 2019). Briefly, at least 1×10^6^ fixed cells were digested using the DpnII restriction enzyme (NEB, R0543M). Chromatin was re-ligated with T4 ligase (Thermo Fisher Scientific), de-crosslinked and precipitated. To check the validity of the experiment, 500 ng of re-ligated DNA were loaded on a 1% gel along with undigested and digested controls. 3C libraries were sheared and adaptors ligated to the libraries according to the manufacturer’s instructions for Illumina sequencing (Agilent). Pre-amplified libraries were hybridized to the custom-designed SureSelect beads (chr13: 54,000,001-57,300,000) (Kragesteen et al. 2018)) and indexed for sequencing (50–100 bp paired-end) following the manufacturer’s instructions (Agilent). Enriched libraries were pooled at 2 nM and loaded for clustering on a Paired-End Illumina Flow cell for an average of 215 mio reads/library. Reads of 100 bases were generated using the TruSeq SBS chemistry on an Illumina HiSeq 4000 sequencer.

## STATISTICAL ANALYSIS AND COMPUTATIONAL ANALYSIS

### ChIP-seq

Single-end reads were mapped to the reference genome NCBI37/mm9 using Bowtie2 version 2.3.4.2 (Langmead and Salzberg 2012), filtered for mapping quality q ≥ 25 and duplicates were removed with SAMtools 1.9. Reads were extended to 250 bp and scaled (1 million/total of unique reads) to produce coverage tracks. BigWig files were visualized in the UCSC genome browser.

### RNA-seq

Single-end reads were mapped to the mm9 reference genome using STAR mapper version 2.5.2a with default settings. Further processing was done according to (Paliou et al. 2019). BigWig files were visualized in the UCSC genome browser. Counting was done using R version 3.6.2 and differential expression was analyzed through the “DEseq2” R package. The DEseq2 R package was also used to produce heatmaps by subtracting from each gene value per condition, given by vst, the mean value of all conditions. Genes were picked according to adjusted p-value, all being significantly differentially expressed between conditions. *Pitx1* fold enrichment between WT GFP+ and GFP-populations was calculated normalizing the total normalized read count per million. Significance in the fold enrichment was calculated using a Student’s t.test (type 2, 1 tail), inputting the normalized counts from each condition (2 normalized counts per genetic background). Expression heatmaps were generated for satellite and mesenchymal markers as defined in Supplementary table S1. For visualization reasons, *Ccr5, Cldn5* and *Col2a1* were added as sub-cluster markers (endothelium immune and condensation) and the forelimb-specific marker *Tbx5* was removed from the marker list. Moreover, genes with expression less or equal to 1 RPKM in all 8 samples (GFP+ wildtype: replicate 1 and 2; GFP-wildtype: replicate 1 and 2, GFP+ mutant: replicate 1 and 2; GFP-mutant: replicate 1 and 2) were removed from the analysis. For the GFP-specific heatmap, we additionally removed all genes with less or equal to 1 RPKM in all 4 GFP-samples. The color of the expression heatmap corresponds to the z-score transformed RPKM values, using the mean and standard deviation per gene based on all 8 samples. Log2FC was calculated by averaging replicates RPKM for each datasets and dividing *Pitx1*^*GFP*^ and *Pitx1*^*GFP;ΔPen*^ values.

### Capture-HiC and virtual 4C

Paired-end reads from sequencing were mapped to the reference genome NCBI37/mm9 using with Bowtie2 version 2.3.4.2 (Langmead and Salzberg 2012) and further filtered and deduplicated using HiCUP version 0.6.1. When replicates were available, these were pooled through catenation (-cat in Python 2.7.11) before HiCUP analysis. Valid and unique di-tags were filtered and further processed with Juicer tools version 1.9.9 to produce binned contact maps from valid read pairs with MAPQ ≥ 30 and maps were normalized using Knights and Ruiz matrix balancing, considering only the genomic region chr13: 54,000,001-57,300,000 (Knight and Ruiz 2013; Wingett et al. 2015; Durand et al. 2016). After KR normalization, maps were exported at 5kb resolution. Subtraction maps were produced from the KR normalized maps and scaled together across their subdiagonals. C-HiC maps were visualized as heatmaps, where contacts above the 99^th^percentile were truncated for visualization purposes. Further details about data processing can be accessed at (Kragesteen et al. 2018). Virtual 4C profiles were generated from the filtered hicup.bam files used also for Capture-HiC analysis. The viewpoint for the *Pitx1* promoter was set at coordinates chr13:55930001-55940000 (10kb bin) and contact analysis was performed over the entire genomic region considered for Capture-HiC (chr13: 54,000,001-57,300,000). A contact pair is considered when one interaction fragment is in the viewpoint and its pair mate is outside of it. The interaction profile was smoothed by averaging over 5kb intervals and was produced as a bedgraph file.

## SINGLE-CELL DATA ANALYSIS

### Processing of sequenced reads

Demultiplexing, alignment, filtering barcode and UMI counting was performed with 10x Genomics Cell Ranger software (version 3.0.2) following manufacture’s recommendations, default settings and mm10 reference genome (version 3.0.0, provided by 10X Genomics, downloaded in 2019). Cell Ranger outputs files for each dataset were processed using the velocyto run10x shortcut from velocyto.py tool (La Manno et al. 2018) (version 0.17.17) to generate a loom file for each sample, using as reference genome the one provided by 10X Genomics and the UCSC genome browser repeat masker .gtf file, to mask expressed repetitive elements. Each loom matrix, containing spliced/unspliced/ambiguous reads, was individually imported in R (version 3.6.2) with the Read Velocity function from the Seurat Wrappers package (version 0.2.0). In parallel, feature filtered output matrices obtained from Cell Ranger were individually loaded into R through the Read10X function of the Seurat package (version 3.2.0, (Stuart et al. 2019). Then, we combined the spliced, unspliced, ambiguous and RNA feature data in a single matrix for each dataset. Subsequently each matrix was transformed into a Seurat object using Seurat package. Therefore, for each sample we obtained for each sample a single Seurat object comprehend by four assays, three of them (spliced, unspliced and ambiguous) were used for downstream RNA velocities estimations and the RNA feature assay was used for downstream gene expression analysis between the samples, as described below.

### Quality control and filtering

Quality control and pre-processing of each Seurat object of our eight samples was performed attending to the following criteria. Cells expressing less than 200 genes were excluded. Additionally, we calculated the reads that mapped to the mitochondrial genome and we filtered out the cells with a mitochondrial content higher than 15%, since high levels of mitochondrial mRNA has been associated to death cells. Also, we excluded cells with a mitochondrial content lower than 1%, since we observed that belongs, in our datasets, to blood cells probably coming from the dissection protocol.

### Individual dataset normalization, scaling and dimensional reduction

After filtering, one by one we normalized the eight datasets following the default Seurat parameters for the LogNormalize method and applying it only to the RNA features assay. We next scaled it by applying a linear transformation and we calculated the most variable features individually for downstream analysis, using standard Seurat parameters. Scaled data was then used for principal component analysis (PCA), we used the 50 PCs established by default, and non-linear dimensional reduction by Uniform Manifold Approximation Projection (UMAP (Leland McInnes et al. 2018)), we used 1:50 dims as input.

### Cell Doublet identification

Pre-process and normalized datasets were individually screened for detection of putative doublet cells. Doublets in each dataset were also excluded using DoubletFinder R package (version 2.0.2) (McGinnis et al. 2019) as described in https://github.com/chris-mcginnis-ucsf/DoubletFinder. The doublet rate (nExp parameter) used was estimated from the number of cells captured and it is as follows : HLWT replicate 1, nExp = 106; HLWT replicate 2, nExp = 123; FLWT replicate 1, nExp = 97; FLWT replicate 2, nExp = 116; HLPitx1^-/-^ replicate 1, nExp = 104; HLPitx1^-/-^ replicate 2, nExp = 122; HL^Pen-/Pen-^ replicate 1, nExp = 118; HL^Pen-/Pen-^ replicate 2, nExp =116. The pK parameter was calculated following the strategy defined by (McGinnis et al. 2019) and is as follow: HLWT replicate 1, pK = 0.12; HLWT replicate 2, pK = 0.005; FLWT replicate 1, pK = 0.09; FLWT replicate 2, pK = 0.04; HLPitx1^-/-^ replicate 1, pK = 0.04; HLPitx1^-/-^ replicate 2, pK = 0.01; HL^Pen-/Pen-^ replicate 1, pK = 0.005; HL^Pen-/Pen-^ replicate 2, pK = 0.005. After filtering, we kept for downstream analysis the following number of cells for each dataset: HLWT replicate 1, 4143 cells; HLWT replicate 2, 4816 cells; FLWT replicate 1, 3802 cells; FLWT replicate 2, 4521 cells; HLPitx1^-/-^ replicate 1, 4049 cells; HLPitx1^-/-^ replicate 2, 4745cells; HL^Pen-/Pen-^ replicate 1, 4600 cells; HL^Pen-/Pen-^ replicate 2, 4518 cells.

### Merge of all datasets and normalization

Once each dataset was individually filtered and doublets were removed, all datasets were merged in a unique Seurat object without performing integration to execute an ensemble downstream analysis of the eight datasets. No batch effect was observed later on in this merged dataset. Subsequently, we normalized our new and unique Seurat object applying the SCTransform normalization protocol (Hafemeister and Satija 2019), with default parameters, over the spliced assay.

### Cell-cycle scoring and regression

Since from the individual analysis of our dataset we observed a part of the variance was explained by cell-cycle genes, we examine cell-cycle variation in the merged dataset. To do so, we assigned to each cell a score based on its expression of a pre-determined list of cell cycle gene markers, following the strategy defined by (Tirosh et al. 2016) and by applying CellCycleScoring function implemented in Seurat. Subsequently, the evaluation of this results, we decided to regress out the cell-cycle heterogeneity. Therefore, we applied to our merged object the SCTransform normalization method, using the spliced assay as source, and adding to the default settings the cell-cycle calculated scores (S.Score and G2M.Scores) as variables to regressed.

### Clustering

After cell-cycle regression, cells were clustered using standard steps of the SCTransform Seurat workflow. Briefly, PCA (npcs = 50), UMAP (dims = 1:50) and nearest neighbors of each cell were calculated. Clusters were determined using Seurat FindClusters function with default parameters and a resolution of 0.2, in that way 10 clusters were defined. Identification of clusters identity was done by calculating the expression difference of each gene between each cluster and the rest of the clusters using the FindConservedMarkers function. We applied this function to each cluster (ident.1) using default parameters, only.pos = TRUE and setting as grouping variable the limb identity of the datasets, in that way we obtained a list of markers for each cluster independent of the limb sample. Clusters with similar marker were combined, therefore we finally worked with 5 clusters (**Fig 1B**): the mesenchyme (that contains 5 out of the 10 clusters), the epithelium (formed by 2 out of 10), and the immune cell cluster, the muscle and the endothelium clusters (composed by only 1 cluster each). We confirmed the expected identity markers were present in the new clustering by running the FindMarkers function with the following parameters logfc.threshold = 0.7; pseudocount.use = 0; only.pos = TRUE; min.diff.pct = 0.15 and all other default parameters (Supplementary Table S1).

### Subsetting and Re-clustering

Since the interest of this work was focus on the populations that in a wildtype hindlimb express *Pitx1* (**Fig 1C**), we subsetted the mesenchyme cluster. To have a better insight on the different cell-types that integrate it, we re-cluster the mesenchyme cluster. To do so, UMAP embedding was calculated with the following parameters: dims = c(1:10), n.neighbors = 15L, min.dist = 0.01, metric = “cosine”, spread = 0.5, all other parameters were default. Cluster resolution after finding neighbors was established at 0.4 to reveal subpopulations. We observed 9 mesenchyme subpopulations (**Fig. 3A**) that we named according to their identity genes. Identity markers were found using FindMarkers on the RNA assay, setting logfc.threshold = 0.3, pseudocount = 0, min.diff.pct = 0.1, only.pos = TRUE and all other parameters as default (Supplementary Table S1).

### RNA-velocity analysis

To perform the RNA velocity analysis on the hindlimb wildtype samples we subset the cells belonging to the 2 hindlimb wildtype replicates. This subsetted Seurat object was saved as h5Seurat file using SeuratDisk package (version 0.0.0.9013) and exported to be used as input of Scvelo (version 0.2.2) (https://www.nature.com/articles/s41587-020-0591-3) in Python (version 3.7.3). Then the standard protocol described in <u>scvelo</u> was followed. Standard parameters were used except npcs = 10 and n.neighbors = 15, to be the same that we used for the UMAP embedding in Seurat.

### Differential Proportion analysis

Statistical differential proportion analysis, to study the differences in clusters cell proportions between the different limb-type conditions, was performed in R using the source code published by (Farbehi et al. 2019) after generating the proportion tables in R. Null distribution was calculated using n= 100,000 and p = 0.1 as in the original reference. Pairwise comparisons were performed between the different condition tested.

### Proximal and distal cell classification

Proximal, distal or NR attribute was given to each cluster based on its Shox2 and Hoxd13 expression. Therefore, ICT, TP, PPP and PC clusters were classified as proximal clusters, DP, DPP, EDC and LDC as distal ones. Meanwhile, Ms cluster that express both markers were not classify to any of them. This classification was added to the Seurat object metadata and used in downstream analysis.

### Pitx1 density plot and cell classification by Pitx1 expression

*Pitx1* normalized expression values, from the RNA assay of the all dataset merged Seurat Object, were extracted in a data frame. This data frame was used to create a density plot using ggplot2 package (version 3.3.2). From the overlay of *Pitx1* density distributions in the HLWT samples and the HL^Pen-/Pen-^ we define the intersection point of 0.3 to classify cells in non/low-expressing and expressing cells. The second intersection point of 1.45 that subclassify these expressing cells in mild- and high-expressing cells was established based on the intersection of the HLWT proximal and distal cells (**Fig 2F**). Therefore, we classified as non/low-expressing cells those with Pitx1 expression values <0.3, as mild-expressing those with Pitx1 expressing values between >0.3,<1.45 and as hig-expressing cells those >1.45. This classification and *Pitx1* expression values were added as new columns to the Seurat object metadata and used in downstream analysis.

## Acknowledgments

We thank Mylène Docquier, Brice Petit and Christelle Barraclough from the iGE3 sequencing facility. We thank Jean-Pierre Aubry, Grégory Schneiter and Cécile Gameiro from the Flow Cytrometry facility. We thank Nicolas Liaudet from the imaging facility and Stéphane Pàges and Laura Batti from Advanced Light Sheet Imaging Center (ALICe). We thank Leon Van Gurp, Sigmar Stricker, Pierre Fabre, Quentin Lo Giudice, Lucille Delisle and Denis Duboule for discussions. We thank Michael Robson, Anna Ramisch, Simon Braun and Christina Paliou for critical reading of the manuscript. We thank all lab members for discussions and critical reading of the manuscript. This study was supported by grants from the Swiss National Science Foundation (PP00P3_176802) and from the Boninchi Foundation.

## Data accessibility

Sequencing data have been deposited at the GEO repository and will be available upon publication.

## Author Contributions

G.A., O.B. and R.R. conceived the project. R.R., O.B., R.P. and A.R. performed the ESC targeting and prepared the cells for aggregation. O.B. performed the Capture-HiC, ChIP-eq and RNA-seq preparations and analyses. R.R. performed the scRNA-seq preparation and analyses. G.A., R.R., and O.B. wrote the manuscript with input from the remaining authors.

---

**Figure S1:**
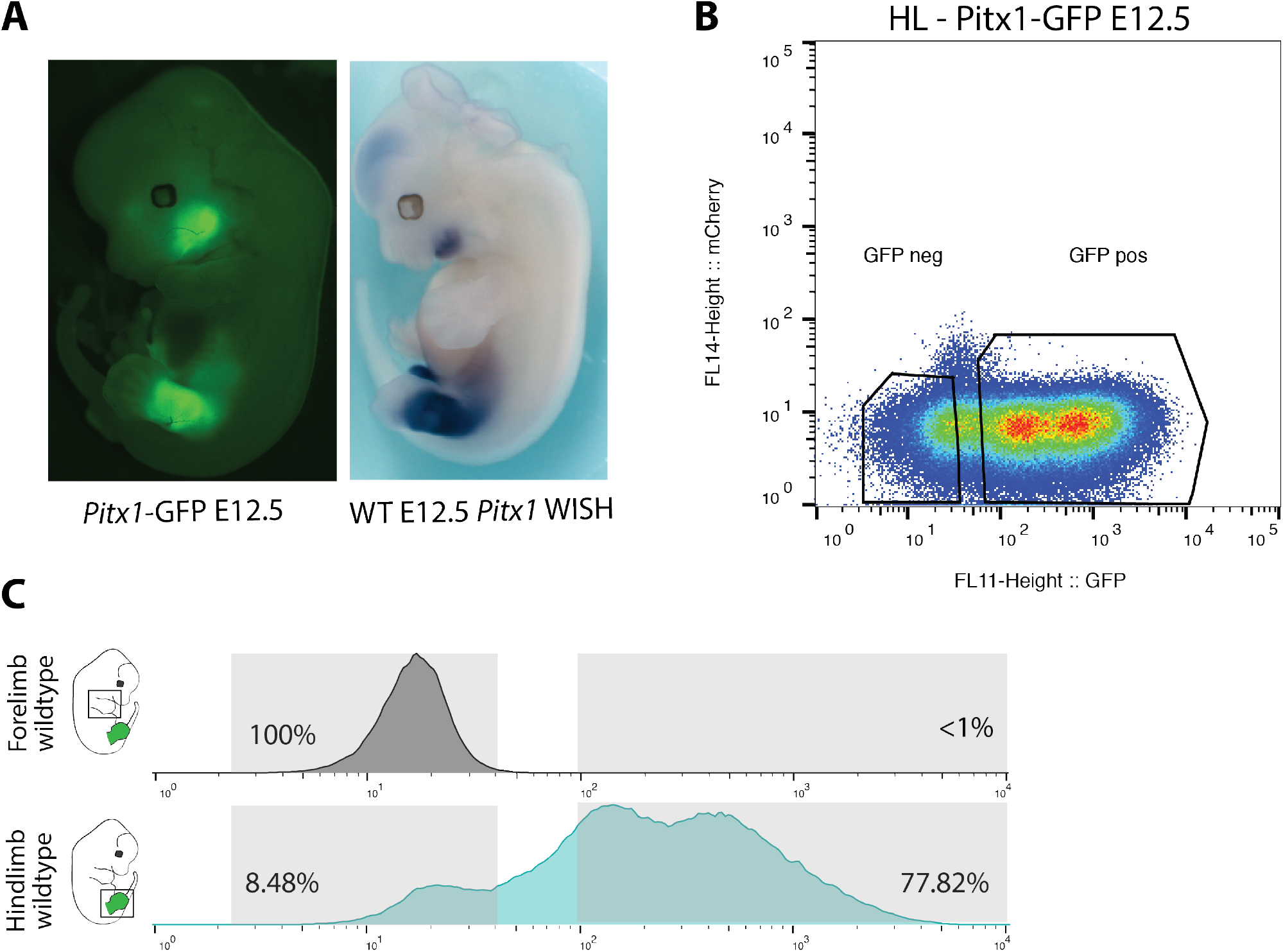
FACS sorting of *Pitx1*^*GFPs/GFPs*^ EGFP cells. **A**. Fluorescence and *Pitx1* WISH of an E12.5 *Pitx1*^*GFP*^ embryo. **B**. Overview of FACS gating with two fluorescent markers (mCherry on the y-axis and EGFP on the x-axis). **C**. *Pitx1*^*GFPs/GFPs*^ forelimb cells were used to delimit the gating of GFP-cells.

**Figure S2:**
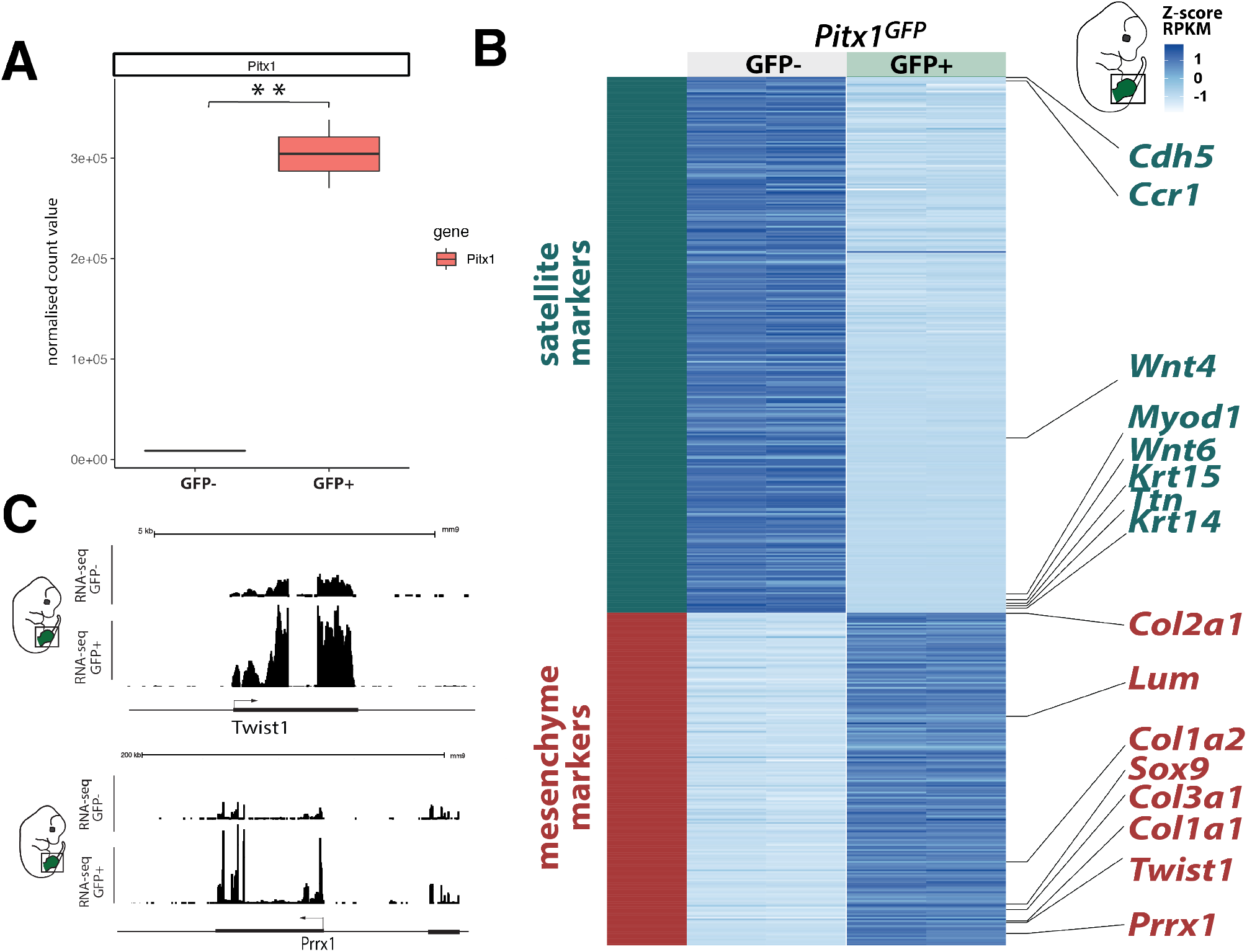
**A**. *Pitx1* normalized RNA-seq counts in GFP- and GFP+ *Pitx1*^*GFP*^ hindlimb cells. ** indicate a p-value <0.01 as calculated with a student t-test. **B**. Heatmap of differentially mesenchymal (red) and satellite (darkgreen) marker genes in GFP+ and GFP-cells in *Pitx1*^*GFP*^ hindlimbs. Note that GFP+ cells are enriched for condensating cells (*Sox9,Col2a1)*, connective tissue (*Col1a1, Col2a1,Col3a1*) and mesenchymal patterning genes (Shox2, *Twist1, Prrx1*). GFP-cells are enriched for muscle (*Myod1, Ttn*), epithelial (*Wnt6*, Wnt4, *Krt14, Krt5*), immune (*Ccr1*) and endothelial cells (*Cdh5*). **C**. RNA-seq tracks at the *Twist1* and *Prrx1* mesenchymal marker loci in GFP- and GFP+ *Pitx1*^*GFP*^ hindlimb cells. Note that in GFP-cells there is expression of these genes, indicating that some GFP-cells are of mesenchymal origin.

**Figure S3:**
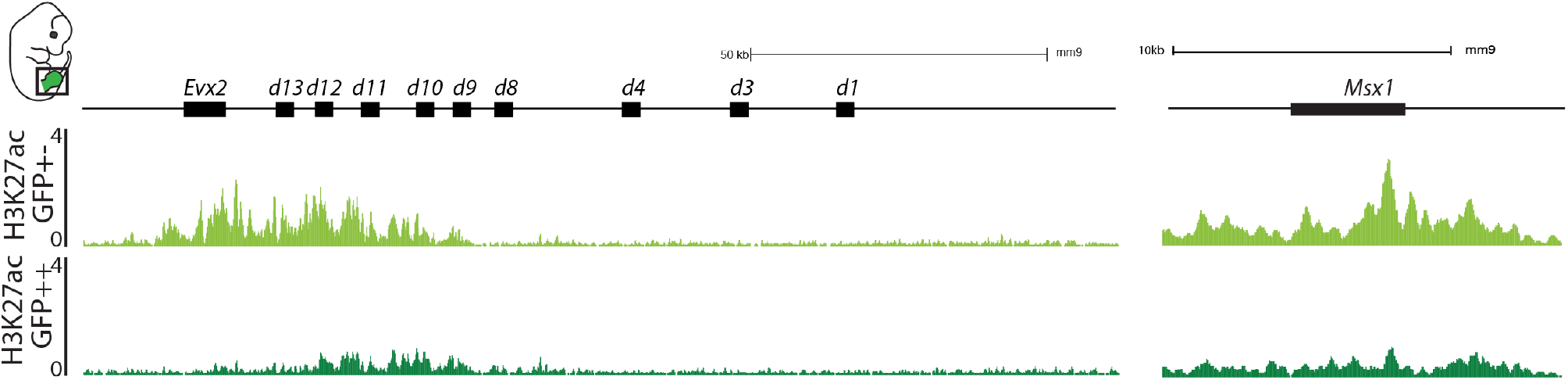
H3K27ac ChIP-seq in mild- and high-expressing hindlimb cell population at the *HoxA* cluster and at the *Msx1* locus. Note the stronger activity in the mildly active cells.

**Figure S4:**
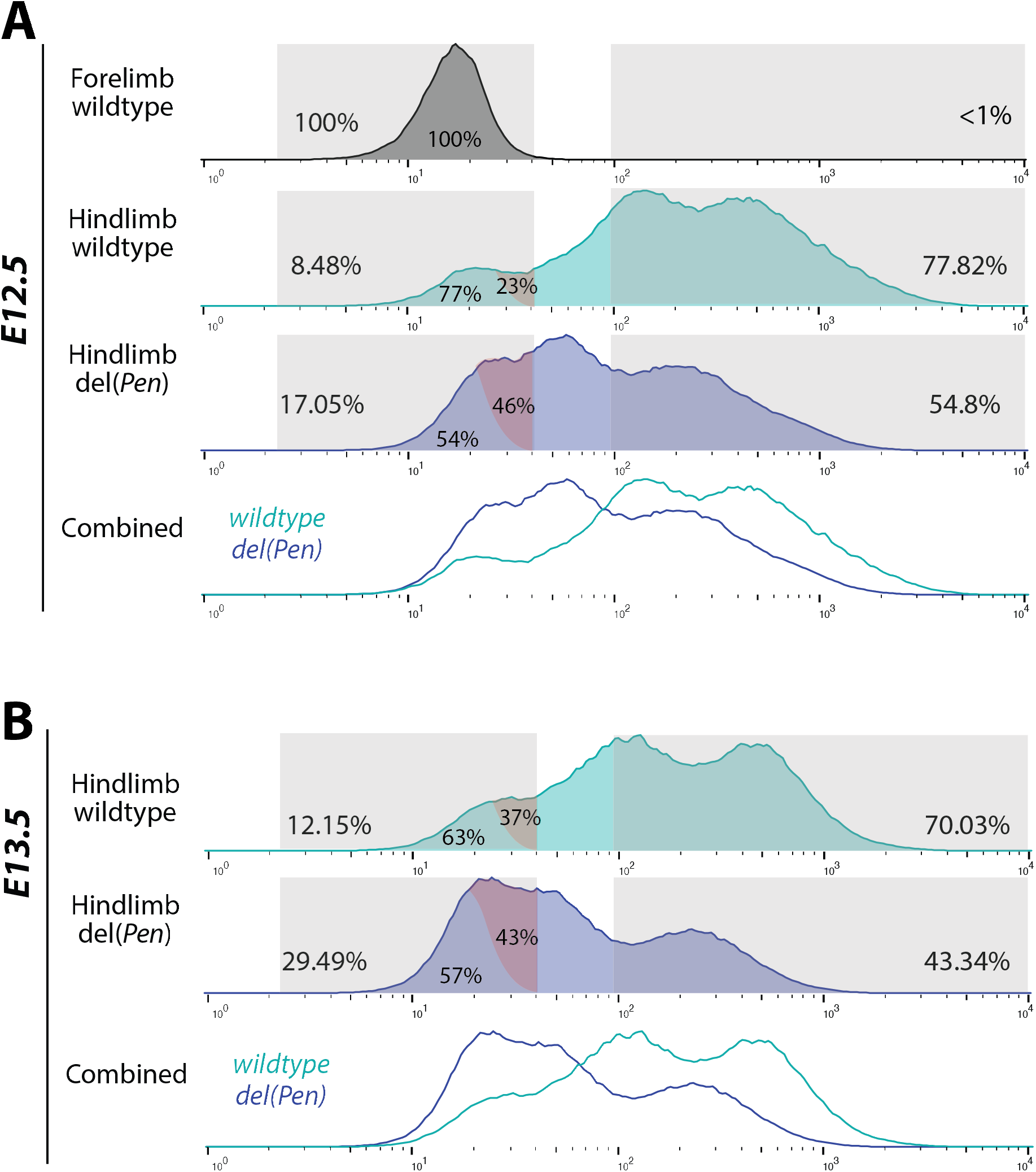
**A**. Comparison of FACS profile in wildtype E12.5 fore- and hindlimb and in del(Pen) hindlimb. Note the increase in the negative cell fraction that includes an increase in non- and low cells. In the negative fraction, the low-expressing cells (red shadowed) increased their proportion with respect to non-expressing cells from 23% in wildtype to 46% in mutant (according to surface ratio). **B**. Comparison of FACS profile in wildtype and del(Pen) E13.5 hindlimbs. In the negative fraction, the low-expressing cells (red shadowed) increased their proportion with respect to non-expressing from 37% in wildtype to 43% in mutant (according to surface ratio).

**Figure S5:**
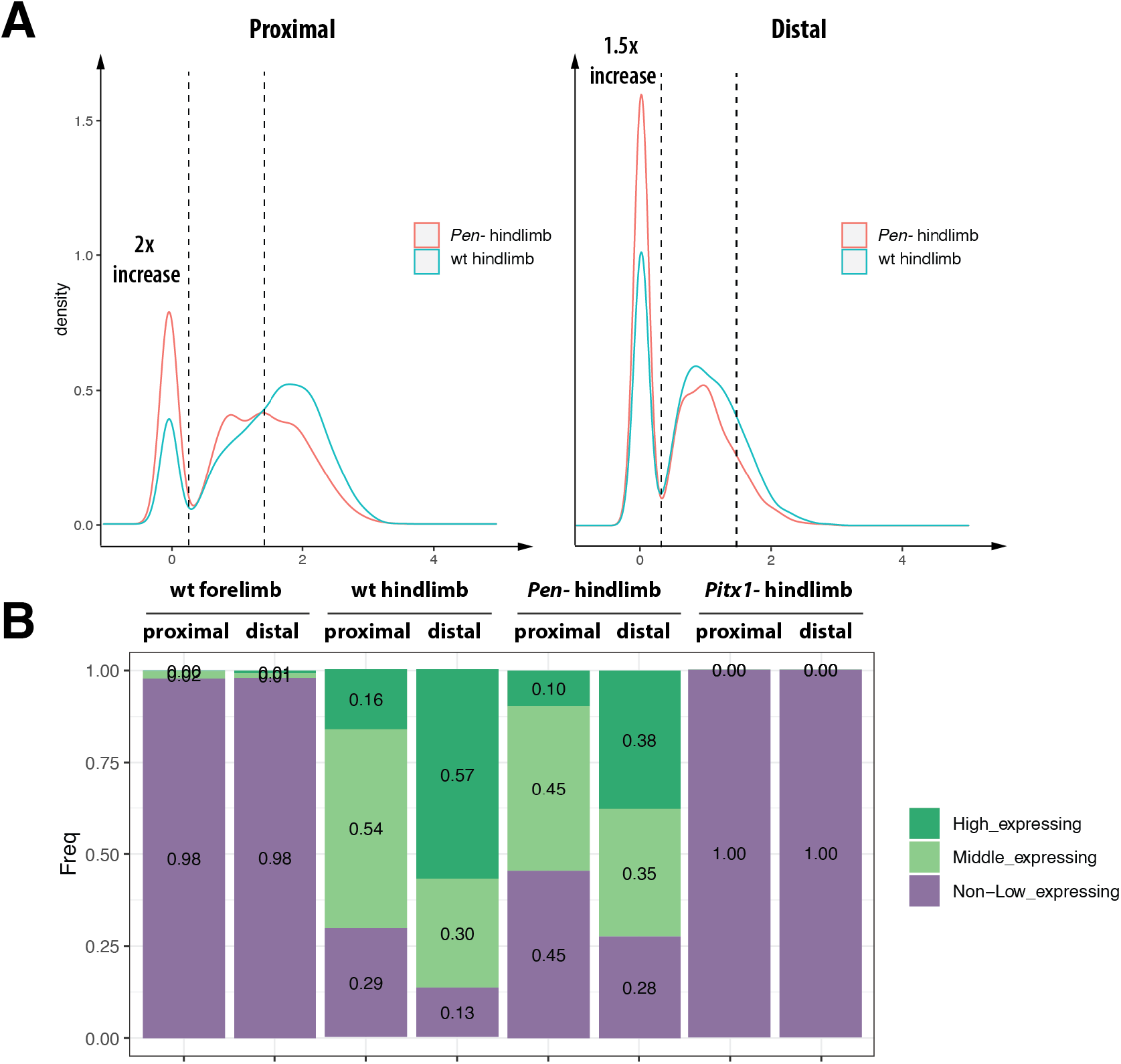
**A**. Distribution of *Pitx1* expression in proximal and distal cells of the hindlimbs in wt and *Pen-*. Note the strong increase in proximal none/low-expressing cell fraction. **B**. Proportion of *Non/low-, mild* and *high*-*Pitx1* expressing cells across conditions.

**Figure S6:**
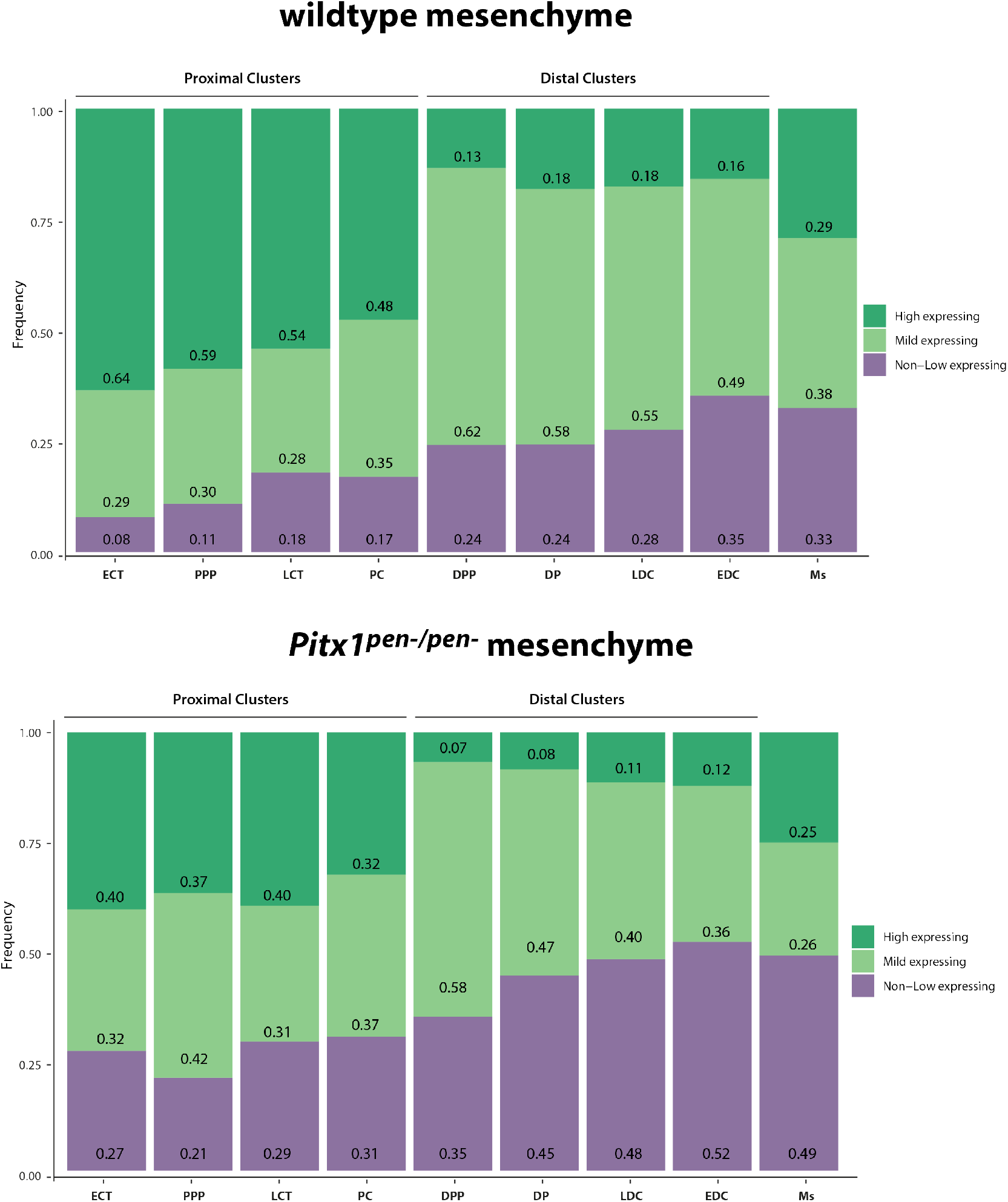
Proportion of *Non/low-, mild* and *high*-*Pitx1* expressing cells in all mesenchymal cell clusters of wildtype and *Pen* deleted hindlimbs.

**Figure S7:**
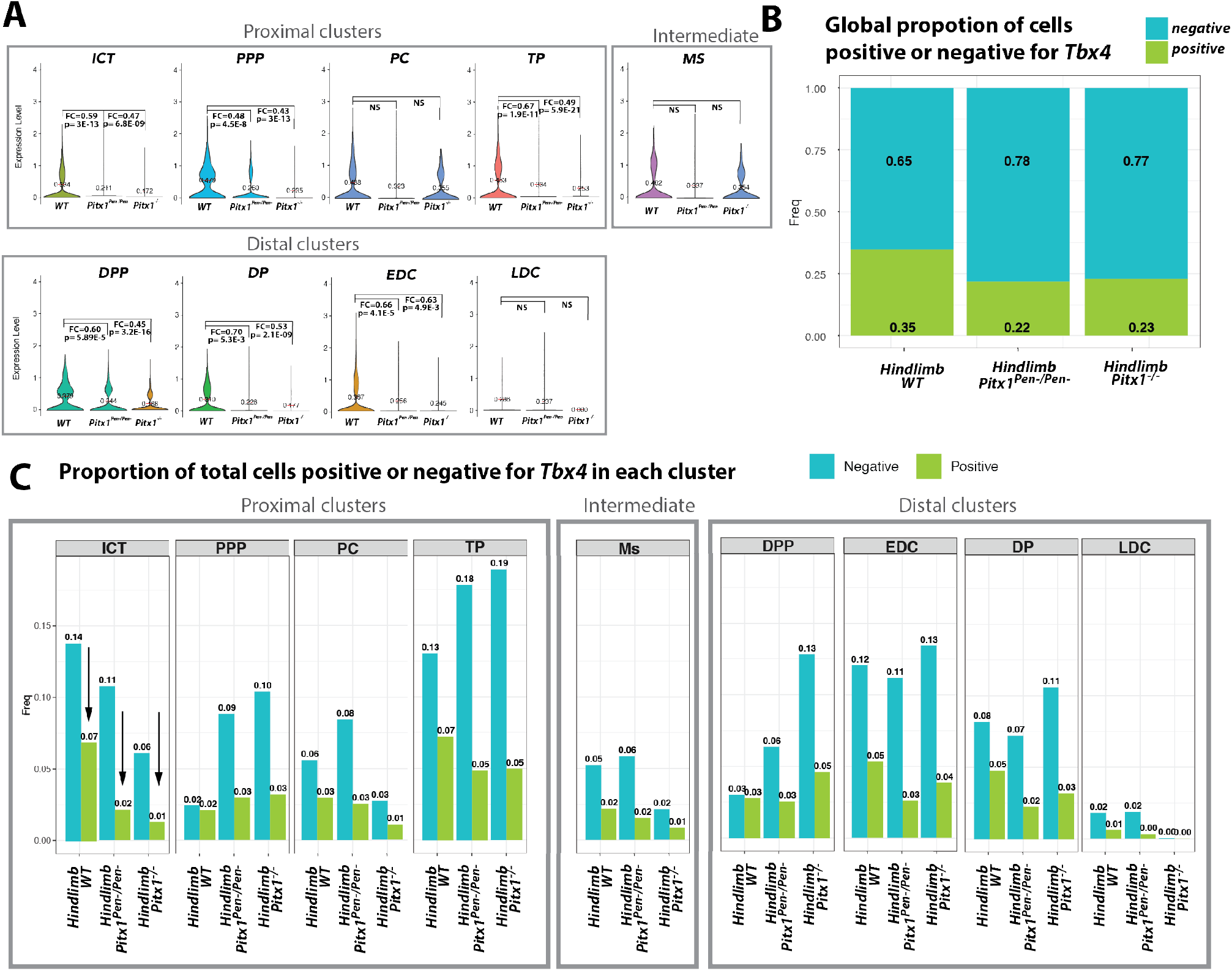
Effect of *Pitx1*^*-/-*^ and *Pitx1*^*Pen-/Pen-*^ on *Tbx4* expression. **A**. Violin plots of *Tbx4* expression in all mesenchymal clusters in wt, *Pitx1*^*-/-*^ and *Pitx1*^*Pen-/Pen-*^ hindlimbs. **B**. Proportion of *Tbx4* expressing and non-expressing cells across the mesenchyme in wt, *Pitx1*^*-/-*^ and *Pitx1*^*Pen-/Pen-*^ hindlimbs. **C**. Proportion of *Tbx4* expressing and non-expressing cells across all mesenchymal clusters in wt, *Pitx1*^*-/-*^ and *Pitx1*^*Pen-/Pen-*^ hindlimbs.

**Figure S8:**
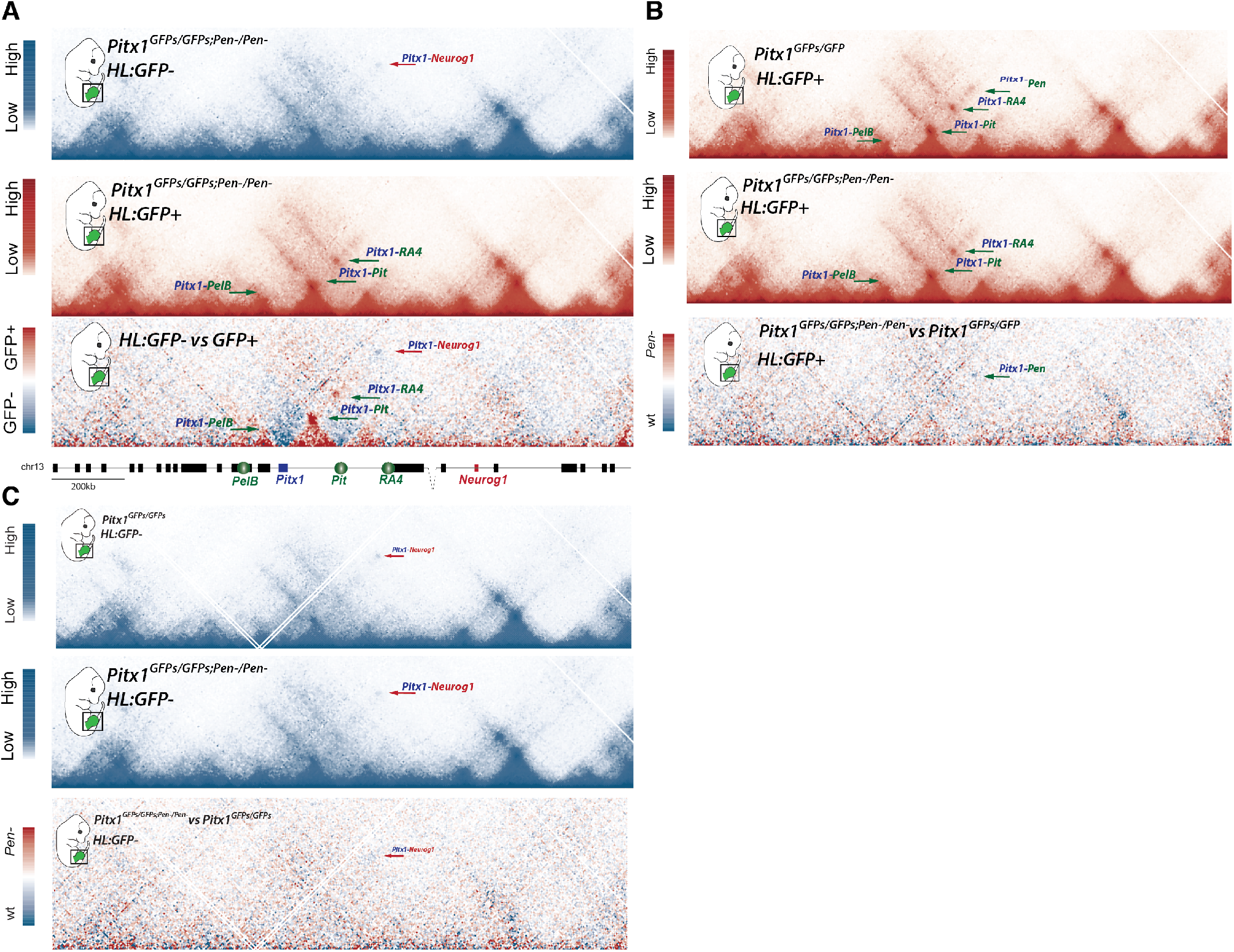
**A**. C-HiC in GFP-(upper panel) and GFP+ (lower panel) sorted hindlimb cells. The lower panel represent a subtraction between the two upper ones. **B**. C-HiC of *Pitx1*^*GFPs*^ GFP+ hindlimb cells (upper panel) and subtraction of this map with the *Pitx1*^*GFP;ΔPen*^ GFP+ cells. **C**. C-HiC of *Pitx1*^*GFP*^ GFP-hindlimb cells (upper panel) and subtraction of this map with the *Pitx1*^*GFP;ΔPen*^ GFP-cells.

## Legends to supplementary tables and video

**Table S1:** Marker genes for single cell clusters

**Table S2:** DeSeq2 analysis of GFP-vs GFP+ *Pitx1*^*GFP*^ hindlimbs. Positive FC indicate enrichment in GFP+ cells and negative FC indicate enrichment in GFP-cells.

**Table S3:** DeSeq2 analysis of *Pitx1*^*GFP*^ vs *Pitx1*^*GFP;ΔPen*^ *GFP-*hindlimb cells. Positive Log2FC indicates enrichment in GFP-cells of *Pitx1*^*GFP;ΔPen*^ hindlimbs and negative Log2FC indicates depletion in GFP-cells from *Pitx1*^*GFP*^ hindlimbs.

**Table S4:** DeSeq2 analysis of *Pitx1*^*GFP*^ vs *Pitx1*^*GFP;ΔPen*^ *GFP+* hindlimb cells. Positive Log2FC indicates enrichment in GFP+ cells of *Pitx1*^*GFP;ΔPen*^ hindlimbs and negative Log2FC indicates depletion in GFP+ cells from *Pitx1*^*GFP*^ hindlimbs.

**Table S5:** sgRNAs used in this study and relative genomic location

**Video S1:** 3D reconstruction of an *Pitx1*^*GFP*^ E12.5 embryo.

